# Antioxidant Response Activating nanoParticles (ARAPas) localize to atherosclerotic plaque and locally activate the Nrf2 pathway

**DOI:** 10.1101/2021.09.16.460323

**Authors:** Sophie Maiocchi, Ana Cartaya, Sydney Thai, Adam Akerman, Edward Bahnson

## Abstract

Atherosclerotic disease is the leading cause of death world-wide with few novel therapies available despite the ongoing health burden. Redox dysfunction is a well-established driver of atherosclerotic progression; however, the clinical translation of redox-based therapies is lacking. One of the challenges facing redox-based therapies is their targeted delivery to cellular domains of redox dysregulation. In the current study, we sought to develop Antioxidant Response Activating nanoParticles (ARAPas), encapsulating redox-based interventions, that exploit macrophage biology and the dysfunctional endothelium in order to selectively accumulate in atherosclerotic plaque. We employed flash nanoprecipitation (FNP) to synthesize bio-compatible polymeric nanoparticles encapsulating the hydrophobic Nrf2 activator drug, CDDO-Methyl (CDDOMe-ARAPas). Nuclear factor erythroid 2-related factor 2 (Nrf2)-activators are a promising class of redox-active drug molecules whereby activation of Nrf2 results in the expression of several antioxidant and cyto-protective enzymes that can be athero-protective. In this study, we characterize the physiochemical properties of CDDOMe-ARAPas as well as confirm their in vitro internalization by murine macrophages. Drug release of CDDOMe was determined by Nrf2-driven GFP fluorescence. Moreover, we show that these CDDOMe-ARAPas exert anti-inflammatory effects in classically activated macrophages. Finally, we show that CDDOMe-ARAPas selectively accumulate in atherosclerotic plaque of two widely-used murine models of atherosclerosis: ApoE^−/−^ and LDLr^−/−^ mice, and are capable of increasing gene expression of Nrf2-transcriptional targets in the atherosclerotic aortic arch. Future work will assess the therapeutic efficacy of intra-plaque Nrf2 activation with CDDOMe-ARAPas to inhibit atherosclerotic plaque progression. Overall, our present studies underline that targeting of atherosclerotic plaque is an effective means to enhance delivery of redox-based interventions.

## Introduction

According to the World Health Organization, cardiovascular disease (CVD) is the most common cause of death in the entire world^1, 2^. A common underlying cause of clinical events is a chronic, long-term process called atherosclerosis, which involves progressive changes to arterial structure and function^3^. Atherosclerosis is characterized by focal lesions formed in the sub-intimal space of large and mid-sized arteries, composed of lipids, fibrotic tissue and inflammatory cells that result in narrowing of the blood vessel lumen^3^. Atherosclerotic lesion rupture and thrombosis leads to embolism that manifests as acute coronary syndrome, myocardial infarction or stroke. A significant early event occurring at vascular sites of disturbed blood flow is endothelial dysfunction, which permits the increased infiltration of activated immune cells, such as monocytes, and low-density lipoprotein (LDL) into the sub-intimal space, where LDL is susceptible to oxidative modification. Infiltrating monocytes differentiate into macrophages, which recognize modified LDL, by surface scavenger receptors. This results in the excessive accumulation of lipids in macrophages, which become foam cells, the hallmark of early fatty-streaks^4^. Ongoing chronic inflammation within the intima results in the eventual formation of more complex atherosclerotic plaques, consisting of an overlying fibrous cap, composed of collagen and smooth muscle cells (SMCs), and a necrotic core, derived from dying foam cells, calcium deposits and cholesterol crystals^5, 6^. Overall, both vascular inflammation and dysregulated redox signalling play important roles in the development of endothelial dysfunction, the progression of atherosclerosis and its clinical manifestations^7, 8^. Although a considerable body of pre-clinical studies indicate that atherosclerosis is driven by oxidative processes^7, 8^, the clinical translation of redox-based therapies is lacking. One challenge related to therapeutically targeting redox dysregulation in atherosclerosis, is the specific delivery of redox-based interventions to cellular domains where redox dysregulation is occurring^8^. Targeted nanomedicine is a promising, and under-studied approach to counter this challenge in CVD^9–13^. Encapsulation of pharmacological therapeutics into nanoparticles can endow them with the capacity to exploit the enhanced permeability retention effect present in atherosclerotic plaque due to gaps in the dysfunctional endothelium^14^, as well as exploit internalization by plaque-resident macrophages^15, 16^.

A promising mechanistic approach to limit oxidative stress and inflammation associated with atherosclerosis is activation of the Kelch-like ECH-associated protein 1/ Nuclear factor erythroid 2-related factor 2 (Keap1/Nrf2) transcription factor pathway^9, 17^. Nrf2 is a master regulator of the cellular response to oxidative or electrophilic stress through the transcription of numerous cytoprotective and antioxidant genes including glutamate-cysteine ligase catalytic subunit (GCLC), NADPH Quinone dehydrogenase (NQO1), heme oxygenase 1 (HO1), and superoxide dismutase 1 (SOD1). Under homeostatic conditions, Keap1 binds Nrf2 in the cytoplasmic compartment and promotes its ubiquitination and downstream proteasomal degradation. However, upon oxidative or electrophilic stress, specific Keap1 cysteine resides are modified, resulting in nuclear translocation of newly generated Nrf2 where it binds to the Antioxidant Response Element (ARE) and/or the Electrophilic Response Element (EPRE) and drives expression of its target cytoprotective genes. Nrf2 and its target genes have local anti-atherogenic effects in the vascular wall, including endothelial cells, vascular smooth muscle cells, and macrophages^18–21^. Nrf2 also limits inflammation by directly impeding transcription of pro-inflammatory cytokines such as IL-1β and Il-6^20^. Moreover, compounds known as Nrf2-inducers (tBHQ^22^, Ebselen^23^, CDDO-Me analogue dh404^24^, and oleanolic acid^25^) augmented endogenous antioxidant systems and limited inflammation to prevent atherosclerosis development or progression in diabetes-aggravated atherosclerosis. One potent nanomolar Nrf2 activator of interest is CDDO-Me, a synthetic triterpenoid analogue of oleanolic acid, which is currently under-going clinical trials^26^. Herein we sought to generate Antioxidant Response Activating Particles (ARAPas), encapsulating the Nrf2-actvator (CDDOMe), for the selective delivery of this Nrf2 activator to atherosclerotic plaque.

## Experimental

Detailed methods of all experiments can be found in the ESI.

### General materials

CDDO-methyl (Sigma Aldrich, St. Louis, MO. SMB00376-100MG), Synperonic-PE-P84 pluronic tri-block co-polymer (Sigma Aldrich, St. Louis, MO. 713538-1Kg), DMSO (Fisher Scientific, D128-1), DMEM (11885-084; Gibco, Grand Island, NY). DMSO (BP231; Thermo-Fisher Scientific, Waltham, MA), F-12 nutrient mix Ham’s media (11765-054; Gibco), Glucose (50-99-7, Sigma-Aldrich), DMEM High glucose (4.5g/L) (11995-065, Gibco), DMEM low glucose (1g/L) (11885092, Gibco), RPMI 1640 media (11875135, Gibco), Heat-inactivated fetal bovine serum (FBS) (16140071; Gibco), Penicillin-Streptomycin 10,000U/mL (15140122, Gibco), Glutamine (200mM, 25030081; Gibco), Paraformaldehyde (158127; Sigma-Aldrich). PBS (20–134; Apex Bioresearch Products, San Diego, CA). Trypsin-EDTA (0.05%) (25300054; Gibco).

### Cell Culture

All cells were cultured in an incubator at 37 °C with 5% CO_2_. RAW 264.7 macrophage cells (ATCC, TIB-71) were cultured in DMEM, high glucose (4.5g/L) supplemented with 10% FBS, 1% Penicillin-streptomycin. H1299 cells containing a GFP fragment retrovirally inserted into the second intron of the NQO1 gene (130207PL1G9) were a gift from Tigist Yibeltal and the Major Lab (UNC, LCCC), Uri Alon and the Kahn Protein Dynamics group^27, 28^. These cells were cultured in RPMI 1640 media supplemented with 10% FBS and 1% Penicillin-streptomycin. CDDO-Me was stored in stock concentrations in DMSO at − 20 °C and diluted directly in media for treatment. Equal volume DMSO was included for all controls. CDDOMe-ARAPas were stored in 1XPBS and diluted into media. Equal volumes of PBS and equal mass concentrations of polymer were included as a control.

### Synthesis of CDDOMe-ARAPas

CDDOMe-ARAPas were synthesized via flash nanoprecipitation (FNP) with a confined impinging jet mixer (CIJ) as previously described^29^. Briefly, CDDO-Me and Synperonic PE-P84 were dissolved in tetrahydrofuran (THF, Fisher Scientific, T425-1) at 5 mg/mL each. PBS was used as the aqueous solvent. The two solvent streams were mixed together with the CIJ into 4 mL of PBS. This suspension was then dialyzed overnight against PBS. To generate fluorescent nanoparticles, DiD (Fisher Scientific, D7757, final concentration 0.25mg/mL) was added to the organic solvent prior to injecting the solvent streams through the CIJ.

### Nrf2 activation assay

H1299 cells were seeded at 20,000 cells per well in a 96 well black glass-bottom plate (Greiner 96 well plates, 655891, VWR). Media was replaced with media containing treatments (1% DMSO, CDDO-Me (10-200nM), CDDOMe-ARAPas (10-200nM) and equivalent concentrations of polymer). Cells were then either imaged continuously over a 40 hr period (intervals of 4 hr), or were imaged once 24 hours following incubation with treatment. Wells were imaged using Gen 5 software (BioTek Instruments, Winooski, VT) on a Cytation 5 plate reader (BioTek Instruments) (37°C, 5% CO2) with a GFP filter cube (BioTek Instruments, part #:1225101) and a Texas Red filter cube (BioTek Instruments: Part # 1225102). Exposure times were fixed for each well. Cells were counted automatically by the Gen 5 software by thresholding in the Texas Red channel.

### Confocal microscopy

Fluorescence images were obtained by using an inverted laser scanning confocal microscope (Zeiss LSM 780; Zeiss, Oberkochen Germany) through a 63x oil immersion objective lens (numerical aperture 1.40, catalog# 420782-9900; Zeiss, Oberkochen, Germany) with a zoom of 2, pixel size: 0.0659×0.0659 µm^2^. Nuclear DAPI was excited by a 405 nm laser diode and images obtained through a detection wavelength of 410-585 nm with a conventional PMT detector. CD11b-Alexa Fluor 555 was excited by an Argon laser (514 nm) and images obtained through a detection wavelength of 524-656 nm with a conventional PMT detector. NP-DiD was excited by a helium-neon laser (633 nm) and images obtained through a detection wavelength of 638-755 nm with a conventional PMT detector. Pinhole size for all channels was set to 1 airy unit. Scanning mode was set to frame. 3D stacks were obtained with dz=1.00 µm. Three dimensional rendering and related surfaces were obtained with IMARIS software v9.7.2 (Bitplane AG, Zürich, Switzerland). Orthogonal views and representative image (slice, z6/13) images were obtained using Fiji (Fiji Is Just ImageJ; NIH).

### Animals and diet

All animal handling and experimental procedures were approved by the Institutional Animal Care and Use Committee at the University of North Carolina – Chapel Hill (IACUC ID: 18-303). 4-6-week-old male ApoE^−/−^ mice (B6.129P2-Apoetm1Unc, stock number: 002052), LDLr^−/−^ (B6.129S7-Ldlrtm1Her/J, stock number: 002207) and C57Bl/6 mice (stock number: 000664) were purchased from Jackson laboratory. C57Bl/6 mice were fed standard chow. Mice were allowed ad libitum access to food and water throughout the study. After 1 week acclimation in the Division of Comparative Medicine (DCM) facility, ApoE^−/−^ and LDLr^−/−^ were placed on a western high-fat diet containing 40% fat, 17% protein, 43% carbohydrate by kcal and 0.15% cholesterol by weight (RD Western Diet, catalog number: D12079Bi). apoE^−/−^ and LDLr^−/−^ mice were fed with the high-fat diet over the course of 8-15 weeks. Mice underwent experimental procedures and were sacrificed when they were approximately either 13-15 weeks old or 20-22 weeks old.

### Statistical analysis

Numerical data are represented as means ± standard deviation. Statistical analyzes were performed using an unpaired Student’s t-test, one-way ANOVA, or factorial ANOVA, or ANCOVA followed by Tukey’s post-hoc test, as appropriate with a p-value < 0.05 considered statistically significant. Multivariate analysis was performed using MANOVA with subsequent univariate analysis and post hoc testing. Statistical analyzes were performed either using OriginLab, Northampton, MA or SPSS, Armonk, NY.

## Results and Discussion

### Generation of CDDOMe-ARAPas

We utilized a novel nanoparticle synthesis technique, flash nanoprecipitation (FNP), to generate polymeric ARAPas encapsulating CDDOMe, a potent nanomolar activator of the Nrf2 transcription factor, as well the lipophilic dye, DiD^29^. FNP allows for the encapsulation of hydrophobic drugs into the core of a protective polymer shell. It is a scalable, continuous synthesis technique which relies upon the rapid mixing of high-velocity streams of liquid, containing the polymers and therapies, in a specially engineered device (confined impinging jet mixer (CIJ)) (**Fig. 1A**). The rapidity of the mixing results in robust generation of homogenous nanoparticles (similar size and therapeutic loading). We chose this technique due to its capacity for high loading efficiency, use of bio-compatible polymers, its scalability, and translational applicability. Indeed, Feng and co-workers have shown that this technique can be scaled up directly from laboratory milligram scale to a 1kg/day nanoparticle production scale, without changes in the size and homogeneity of nanoparticles^30^.

**Figure 1.**
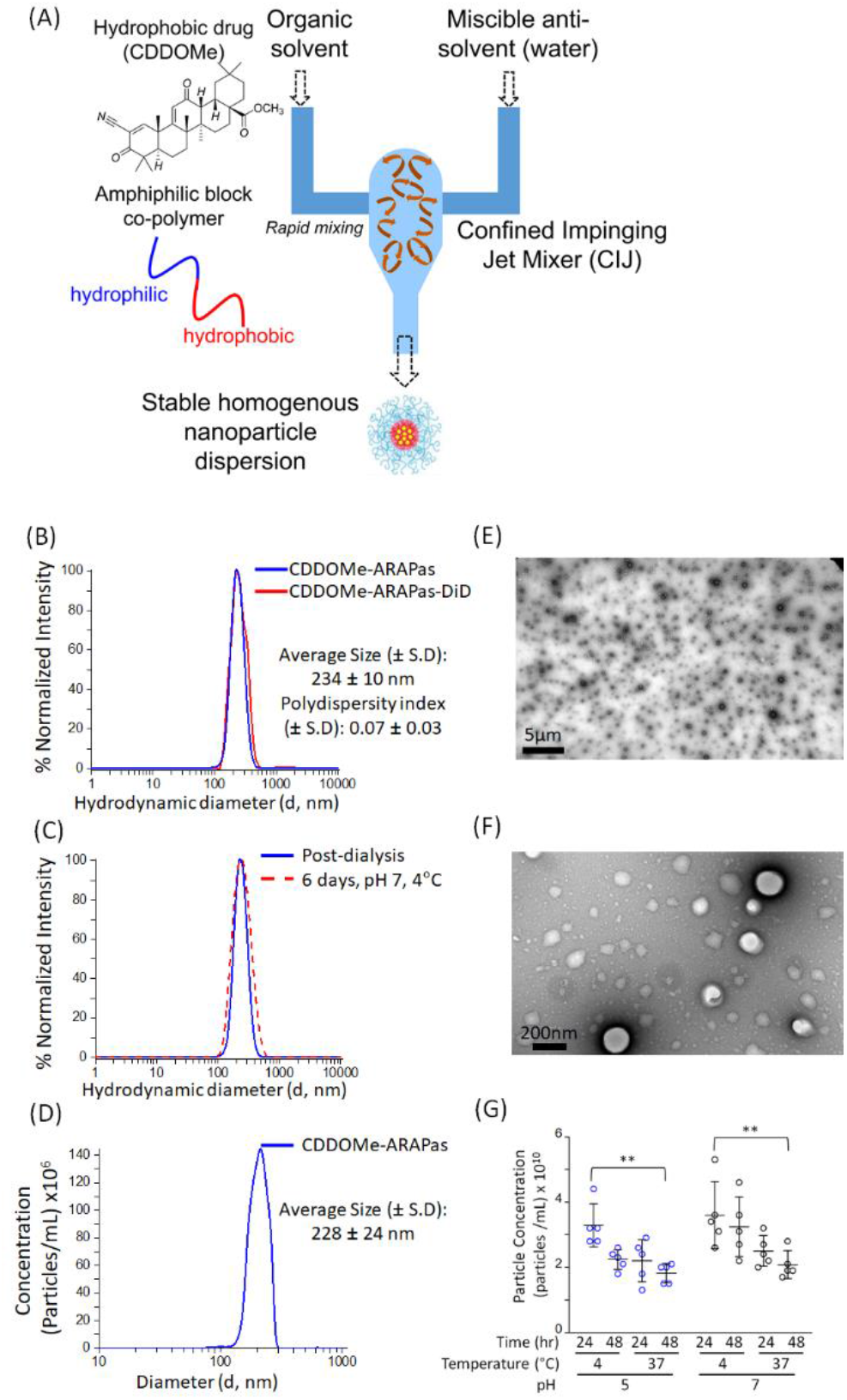
Synthesis and Characterization of CDDOMe-ARAPas. **(A) Schematic overview of the Flash Nanoprecipitation (FNP) method** to generate CDDOMe-ARAPas using a confined impinging jet (CIJ) mixer. **(B) Hydrodynamic diameter and polydispersity index (PDI)** of CDDOMe-ARAPas with or without the lipophilic fluorescent dye, DiD (25μg/mL final concentration). These were measured via an Intensity Distribution with Dynamic Light Scattering (DLS). **(C) Change in hydrodynamic diameter of CDDOMe-ARAPas following 6 days of incubation at 4°C, pH 7.4**. DLS was used to show that the size was unchanged relative to freshly prepared CDDOMe-ARAPas. **(D) Hydrodynamic diameter of CDDOMe-ARAPas determined via nanosight nanotracking analysis**. The data for **(A)-(D)** represent the mean of 3-24 separate preparations of CDDOMe-ARAPas. **(E) TEM of CDDOMe-ARAPas**. Particles were adsorbed onto copper 400 mesh TEM grids were negatively stained with 2% uranyl acetate and imaged via TEM at 5000x magnification (scale bar = 5μm) and at **(F) 100,000x magnification** (scale bar = 200 nm). **(G) Particle concentration as a function of temperature, pH and time**. CDDOMe-ARAPas were diluted 5-fold either into 10mM PBS, pH 7.4 or into 5mM citrate buffer, pH 5 and maintained at either 4°C or 37°C for up to 48 hr. Total particle concentration was assayed at both 24 and 48 hr incubation via Zetaview Nanoparticle Tracking. A 3-way ANOVA to assess the effect of pH, time and temperature was conducted, following by post-hoc Tukey Pairwise comparisons (** P < 0.01).

We generated CDDOMe-ARAPas using the tri-block copolymer synperonic-PE-84 and measured their hydrodynamic diameter by dynamic light scattering (DLS). We determined that they had an average size of 234 ± 10 nm (**Fig. 1B, Table 1**). DLS also revealed a polydispersity index (PDI) of 0.07 ± 0.03, demonstrating that they were highly uniform, which is indicated by a PDI <0.2^31^. To make these nanoparticles fluorescent for visualization in cells and *in vivo*, we included the lipophilic dye, DiD, which resulted in nanoparticles of similar size and polydispersity (**Fig. 1B, Table 1**). Additionally, we found that the nanoparticle size remained unchanged over a period of up to 6 days, when the nanoparticles were stored at 4°C in pH 7.4 10mM PBS (**Fig. 1C**). As an orthogonal measurement, we also performed nanosight nanotracking analysis (NTA) and found a similar average diameter for CDDOMe-ARAPas of 228 ± 24 nm (**Fig. 1D, Table 1**). Moreover, we confirmed a spherical shape and uniform size via transmission electron microscopy (TEM) (**Fig. 1E, F**). Finally, we examined particle stability of CDDOMe-ARAPas, by using total particle concentration as an index, measured with the Zetaview nanoparticle tracking instrument (**Fig. 1G**). CDDOMe-ARAPas were incubated for up to 48 hrs at either 4 or 37°C, and at pH 5 or pH 7.4. We analyzed the effect of temperature, time, and pH upon CDDOMe-ARAPas particle concentration with a factorial ANOVA. The 3-way ANOVA model was significant (F(8,36) = 3.5, P = 0.004). A simple main effects analysis revealed that time (F(1,36) = 5.3, P = 0.03), pH (F(1, 36) = 3.9, P = 0.06) and temperature (F(1, 36) = 15.4, P < 0.0001) all significantly affected particle concentration. However, amongst these variables temperature had the largest effect size ω^2^_p_ = 0.24, with the effect size of pH and time being ω^2^_p_ = 0.06 and ω^2^_p_ = 0.09, respectively. None of the interactions of each of the variables was significant. A post-hoc pairwise comparison with Tukey correction found a significant difference between the particle concentration at 24 hr and 4°C vs 48 hrs and 37°C with particles at either pH 5 or pH 7 (**Fig. 1G**). Moreover, we found that although there is a drop in concentration, this does not relate to particle aggregation as DLS measurements continue to show homogenous populations of the same size (data not shown). To determine the loading efficiency and loading capacity, we performed dialysis to remove unincorporated CDDOMe, and then measured the remaining CDDOMe concentration by HPLC-UV-VIS analysis of the nanoparticles. Loading efficiency was determined by measuring the amount of CDDOMe encapsulated in freshly synthesized nanoparticles vs. those that had been dialyzed overnight. Loading capacity was determined by measuring the mass of CDDOMe present in a solution of solubilized nanoparticles and dividing this by the total maximum mass of CDDOMe and polymer present (i.e. assuming there was no loss of polymer in the nanoparticle generation). In this manner, the loading efficiency was found to be 95.6 ± 5.3 %, with a calculated loading capacity of 32 ± 5 %. FNP is highly useful due to the rapid formulation as well as the high cargo loading^29^. Our studies support this capacity for high loading efficiency resulting in high loading capacity. At the time of writing there appears to be only one other literature report of a polymeric formulation of CDDO-Me with PLGA, where the drug loading capacity was reported to be 2.9 ± 0.2%, after being generated by solvent displacement method^32^.

**Table 1.** Nanoparticle characterization.

| Nanoparticle | Hydrodynamic Diameter (d, nm) | Polydispersity Index (PDI) $\pm 1$ S.D | % Loading Efficiency | Loading Capacity (wt%) |
| --- | --- | --- | --- | --- |
| CDDOMe-ARAPas | $234 \pm 10^a$<br>$228 \pm 28^b$ | $0.07 \pm 0.03$ | $95.6 \pm 5.3$ | $32 \pm 5$ |
| DiD-CDDOMe-ARAPas | $236 \pm 10^a$ | $0.08 \pm 0.04$ | | |

<sup>a</sup>Intensity distribution, dynamic light scattering (DLS)
<sup>b</sup>Nanosight Nanotracking Analysis (NTA)
Values represent the means $\pm 1$ standard deviation of 3-12 independent measurements.

### Time-dependent internalization of CDDOMe-ARAPas by RAW macrophages

Macrophages comprise the bulk of atherosclerotic plaque. The CDDOMe-ARAPas were designed to be close to 200nm as the literature shows optimal macrophage internalization of nanoparticles around this size^15, 16^. Previous literature has also supported that size is an important determinant for accumulation in atherosclerotic lesions, with 200nm-sized liposomes providing the best accumulation^16, 33^. We utilized a murine macrophage cell line, RAW 264.7 macrophages, to examine the association and internalization of CDDOMe-ARAPas with macrophages. CDDOMe-ARAPas were loaded with DiD lipophilic dye and then incubated with RAW 264.7 macrophages over a period of up to 18 hours. Firstly, their association with macrophage cells was followed by kinetic live cell fluorescence imaging (**Fig. 2A,B**), plotting the fluorescence intensity/cell against time. In previous literature, human macrophages internalized 200nm human serum albumin nanoparticles over 4-6 hrs, and then plateaued from 10 hrs, with measurements extending out to 24 hrs^34^. Similarly, in a study by Yu and colleagues^35^ using THP-1 macrophages and studying the internalization of 30, 40 and 100nm polymer-coated iron oxide nanoparticles over 24 hrs, they found that nanoparticle internalization plateaued following 10 hrs, and was described via an exponential function. We also fitted our results to an exponential growth function. The regression was significant (F(3,9) = 11.2, P = 0.004 R^2^ = 0.56), indicating that with time, RFU/cell due to CDDOMe-ARAPas association increases. Our results showed that the nanoparticle internalization remained in a growth phase and had not plateaued between 12 and 18 hrs. Differences in our study that may explain this discrepancy with published literature include that the kinetic assay presented quantifies both nanoparticle external association and internalization, which may not follow the same kinetics as simply internalization. Additionally, we did not follow internalization up to 24 hr, where it may have plateaued from the 18 hr time point. To confirm internalization, we also visualized the CDDOMe-ARAPas, following an 18 hr incubation in RAW macrophages, using confocal microscopy and co-staining for an external cell surface marker, cd11b (**Fig. 2C**). Three-dimensional surface rendering of the confocal fluorescence images indicated that the DiD-CDDOMe-ARAPas were inside the murine macrophages (**Fig. 2D**). To quantify internalization by murine macrophages, we also performed a temperature dependent assay whereby DiD-CDDOMe-ARAPas were incubated with RAW macrophages over 18 hr at either 4 or 37°C (**Fig. 2E**). At 4°C, energy-dependent internalization processes are inhibited, therefore the signal derived from RAW macrophages incubated at this temperature is CDDOMe-ARAPas that are associated with the external cell surface. We analyzed the effect of temperature upon μg DiD/μg cell protein using a t-test. This revealed a significant difference between 4°C vs 37 °C (F(1,4) = 39.7, P = 0.00324), with an effect size of ω^2^ = 0.87, indicating that the majority of signal is due to internalized CDDOMe-ARAPas rather than externally associated nanoparticle. Overall, our studies confirm that murine macrophages significantly internalize CDDOMe-ARAPas in a time-dependent manner.

**Figure 2.**
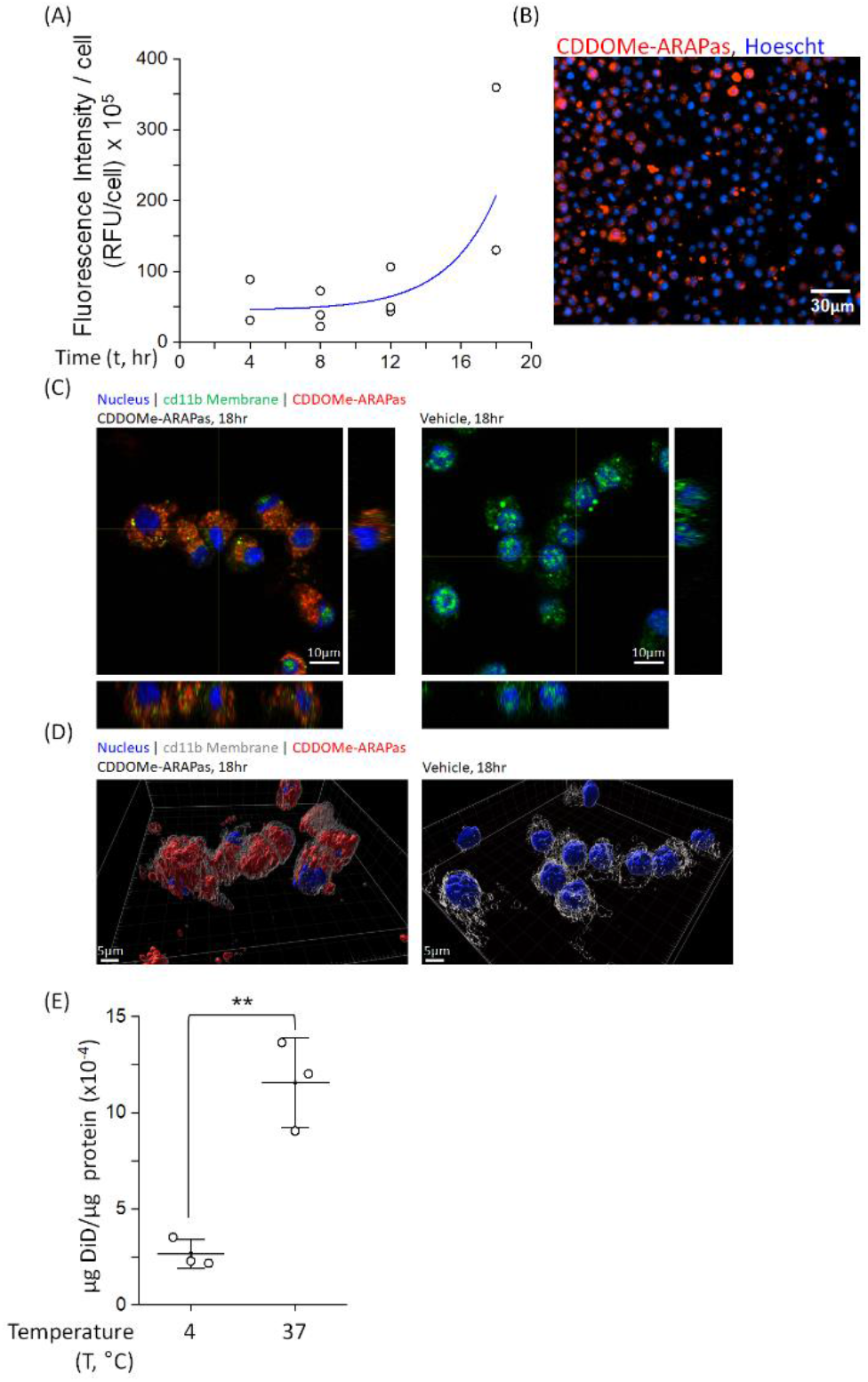
Internalization of CDDOMe-ARAPas by murine macrophages. **(A) Kinetic association of CDDOMe-ARAPas with murine macrophages.** RAW macrophages stained with Hoechst dye (10μg/mL, 30 minutes) were treated with fluorescent DiD-CDDOMe-ARAPas (0.1mg/mL final, 20% PBS) over the course of 18 hrs (37°C, 5% CO2). Fluorescent images of RAW macrophages were taken at 4 hour intervals and thresholded fluorescence area was quantified per cell, an average of 800 cells were counted per well. Data represents N=3 independent experiments in 6-12-tuplicate. **(B) A representative image of macrophage-associated fluorescent DiD-CDDOMe-ARAPas at 18 hr**. Scale bar is 30μm. **(C) Z-projection and orthogonal slices of internalized DiD-CDDO-Me-ARAPas in cd11b stained macrophages**. RAW macrophages were incubated with fluorescent DiD-CDDOMe-ARAPas (0.1mg/mL final, 20% PBS) for 18 hrs (37°C, 5% CO2), and then fixed and counter-stained with cd11b antibody and DAPI. Scale bar is 10μm. **(D) 3D rendered surface image of cd11b membrane, and internalized DiD-CDDO-Me-ARAPas in murine macrophages**. Scale bar is 5μm. **(E) Temperature-dependent internalization of DiD-CDDOMe-ARAPas at 18 hr by RAW macrophages**. Macrophages were incubated with DiD-CDDOMe-ARAPas as described in **(C)-(D)**, and then washed of un-associated ARAPas and the cell lysate assayed for fluorescence intensity and protein content. Data represents means ± 1 S.D. (N=3 independent experiments in triplicate). *, p < 0.05, **, p <0.01.

### Activation of Nrf2 transcription factor by CDDOMe-ARAPas

Various methods exist to determine the release kinetic profile of hydrophobic drugs from nanoparticles^36^, however irrespective of the method, the underlying assumption is that the experiment satisfies sink conditions, where the volume of medium must be at least three times that required to form a saturated solution of the drug. This condition is commonly not satisfied^37, 38^, as for highly insoluble drugs, such as CDDOMe, this can be quite challenging due to large volumes of release medium required and subsequent difficulties analyzing low concentrations of drug. Abouelmagd and co-workers illustrated the issues with two main methods (dialysis and centrifugation) leading to different conclusions of paclitaxel release from nanoparticles^38^. A problem arising from these methods is a mismatch of expected bioactivity either *in vitro* or *in vivo*. Instead, in order to confirm that drug was released by the CDDOMe-ARAPas, we chose to utilize an *in vitro* functional assay, which reflects Nrf2 activation^27, 28^. In this assay, H1299 cells containing a GFP fragment retrovirally inserted into the second intron of the NQO1 gene, a canonical Nrf2-regulated gene^39^, are used to determine Nrf2 activation. Upon Nrf2 activation, GFP fluorescence is observed, which can be quantified as thresholded fluorescence intensity and normalized to cell number due to a Cherry Red nuclear fluorescence. The cells were incubated for 24 hours with increasing doses of either CDDOMe or CDDOMe-ARAPas and then GFP fluorescence intensity/cell measured, of which representative images are shown in **Fig. 3A** and quantified in **Fig. 3B**. To analyze the effect of treatment (CDDOMe-ARAPas and CDDOMe) on Nrf2 activation we conducted an ANCOVA to control for the effect of concentration. The results show that the model fit was significant (F(2,76) = 27, P < 0.0001). Simple main effect analysis showed that dose had a statistically significant effect upon fluorescence intensity/cell ((F(1,76) = 47.5, P < 0.0001)) with an effect size of ω^2^_p_ = 0.37, whilst treatment with either CDDOMe-ARAPas vs CDDOMe alone also had a statistically significant effect upon fluorescence intensity/cell ((F(1,76) = 12.5, P = 0.001)), with an effect size of ω_2_^p^ = 0.13. Thus, we show that that CDDO-Me, a well-known Nrf2 activator, activates the transcription of the canonical down-stream target of Nrf2, NQO1, in a dose-dependent manner (10-400 nM). We additionally show that CDDOMe-ARAPas activate Nrf2, albeit in a significantly different, lesser manner compared to CDDOMe, suggestive of delayed release of CDDOMe from the ARAPas. In previous literature, D’Addio and colleagues indeed reported release of lipophilic compounds from polymeric nanoparticles generated by flash nanoprecipitation^37^, thus our results are consistent with the reported behavior of polymeric nanoparticles generated in a similar manner.

**Figure 3.**
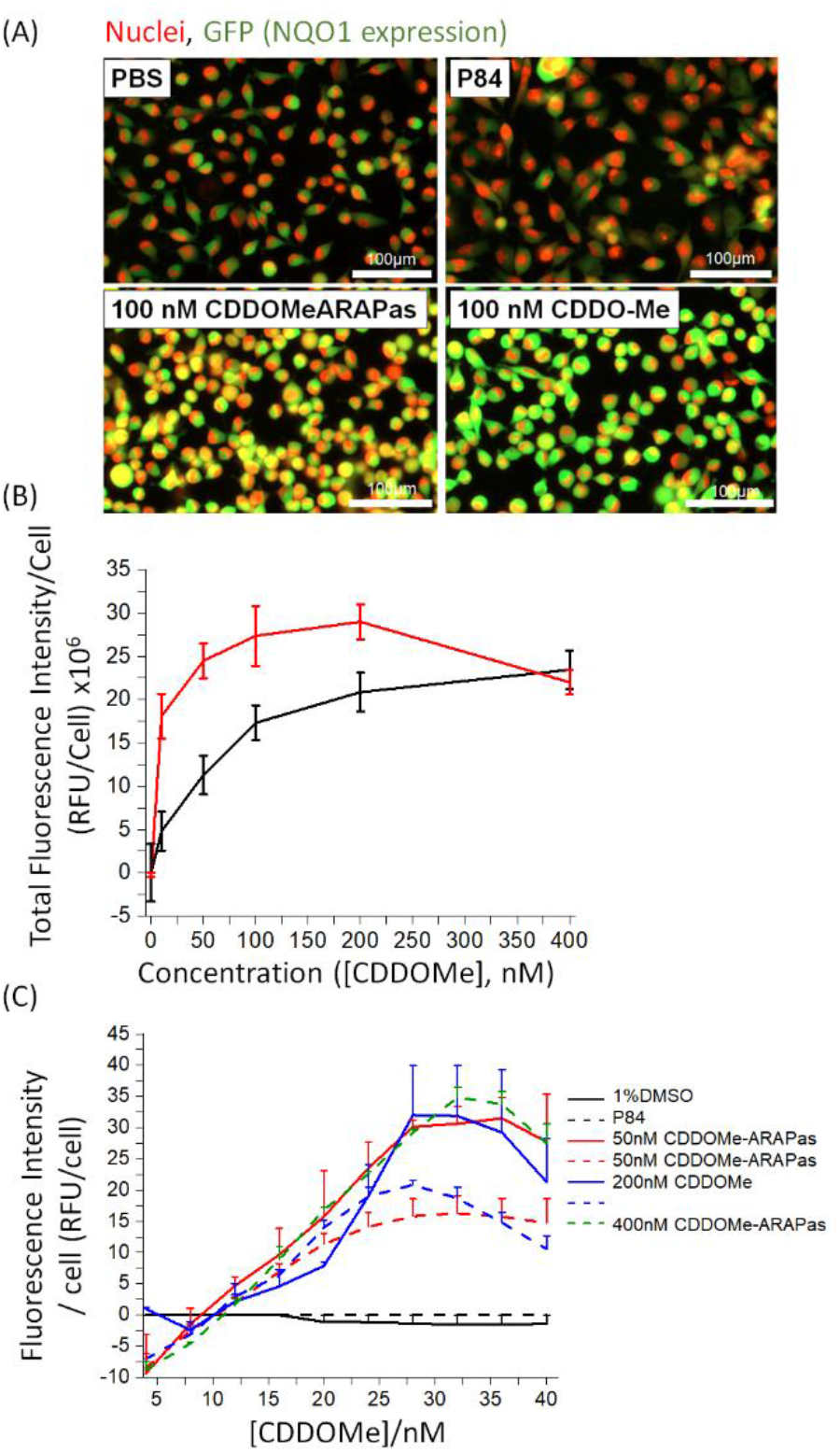
Activation of Nrf2 by CDDOMe-ARAPas. H1299 cells were treated with either 1% DMSO, P84 polymer (at equivalent concentrations as present in most concentrated NP preparations), CDDOMe (10-400nm), or CDDOMe-ARAPas (10-400nM) for 24-40 hours (37°C, 5% CO_2_). **(A) Representative images of live cell GFP-NQO1 fluorescence** as a proxy of Nrf2 activation in H1299 cells after 24 hours as collected by the Cytation 5 plate reader. Scale bar is 100μm. **(B) Dose-dependent activation of NQO1 transcription by CDDOMe and CDDOMe-ARAPas**. GFP fluorescence intensity per cell was quantified with an average of 200 cells counted per measurement. Data represents the means ± 1 S.D. of N=4-5 independent biological experiments, conducted in sextuplicate. **(C) Time-dependent activation of NQO1 transcription by CDDOMe and CDDOMeNPs**. H1299 cells were treated with either 1% DMSO, P84 polymer (at equivalent concentrations as present in most concentrated NP preparations), CDDOMe (solid line, 50-200nm), or CDDOMe-ARAPas (dashed line, 50-400nM) for up to 40 hours. GFP fluorescence intensity per cell was quantified with an average of 200 cells counted per measurement. Data represents the means ± SEM, N=2-4 independent biological experiments.

We then examined the activation of Nrf2 by CDDO-Me and CDDOMe-ARAPas over the course of 40 hours, with kinetic live cell imaging, at varying concentrations (**Fig. 3C**). Herein we found that GFP expression is sustained between 24-40 hr. We additionally recapitulated the trend observed in **Fig. 3B**, wherein higher concentrations of CDDOMe-ARAPas are required (up to 400nM) to sustain GFP expression at similar levels to un-encapsulated CDDO-Me. Taken together, this data indicates firstly, that CDDOMe is released in a delayed manner by CDDOMe-ARAPas and secondly, is capable of activating NRf2 *in vitro*.

### Toxicity of CDDOMe-ARAPas

We also assayed the effect of both CDDOMe and CDDOMe-ARAPas upon macrophage viability with a MTT assay (**Fig. 4A**) and an Annexin V/Apoptosis flow cytometry assay (**Fig. 4B-C**). Naïve macrophages were incubated with increasing concentrations of either CDDOMe or CDDOMe-ARAPas at 0-2000nM over 24 hours and in the case of MTT, then incubated with MTT reagent for a further 4 hours. To determine the EC_50_ we fitted the MTT data with a growth/sigmoidal curve with a dose response function where the upper asymptote was fixed at 100%. The calculated EC_50_ were 274 (95% CI 220 – 328) and 378 (95% CI 348 – 408), for CDDOMe and CDDOMe-ARAPas respectively (**Fig. 4A)**. The lower EC_50_ for CDDOMe compared to CDDOMe-ARAPas supports a slower release of CDDOME from the ARAPas, consistent with data in **Fig. 3**. Analysis of the flow cytometry AnnV/apoptosis data (**Fig 4B-C**) determined that the total viable cell number following either CDDOMe or CDDOMe-ARAPas treatment was reduced in a dose dependent manner. We used multivariate ANOVA (MANOVA) to compare the effect of treatment (CDDOMe or CDDOMe-ARAPas) and concentration on total live, and total dead cell populations (where early and late apoptotic and dead cell populations were grouped together). This revealed a significant difference due to concentration, Wilk’s Lambda λ = 0.302, (F(12,62) = 4.24, P<0.001), with a large effect size estimate ω ^2^_Mult_ = 0.637 on the live cell population. No difference was found when comparing the effect due to treatment with either CDDOMe or CDDOMe-ARAPas (F and p for that main effect). A separate univariate analysis (2-way ANOVA) revealed that the significant effect was upon the live cell population (F(6,32) = 5.72, P < 0.001), but not the dead cell population (F(6,32) = 0.993, P = 0.45). Overall, this indicates that treatment with CDDOMe or CDDOMe-ARAPas significantly reduced the total viable cells without affecting total dead cells (**Fig. 4B-C**). Taken together, these results suggest that lower concentrations of CDDOMe and CDDOMe-ARAPas reduced viable cell number without significant toxicity, indicative of inhibition of proliferation. Previous work by Khoo and co-workers with a related compound, CDDO-imidazole, found that 100nM did not significantly affect viability in HepG2 cells, although 1000 nM led to a loss of ~94% of absorbance in the MTT assay^40^. They also additionally found that 100 nM CDDO-imidazole did not results in toxicity in RAW 264.7 cells. This is in line with our results, albeit in the murine macrophage cell line (RAW 264.7 cells), and an analog of this chemical. In support of the suggestion that CDDOMe may affect macrophage proliferation, in another study, Probst and colleagues^41^ found that an analog of CDDO-Me inhibited growth in 8 different cancer cell lines with an IC_50_ ranging from 159 to 363nM.

**Figure 4.**
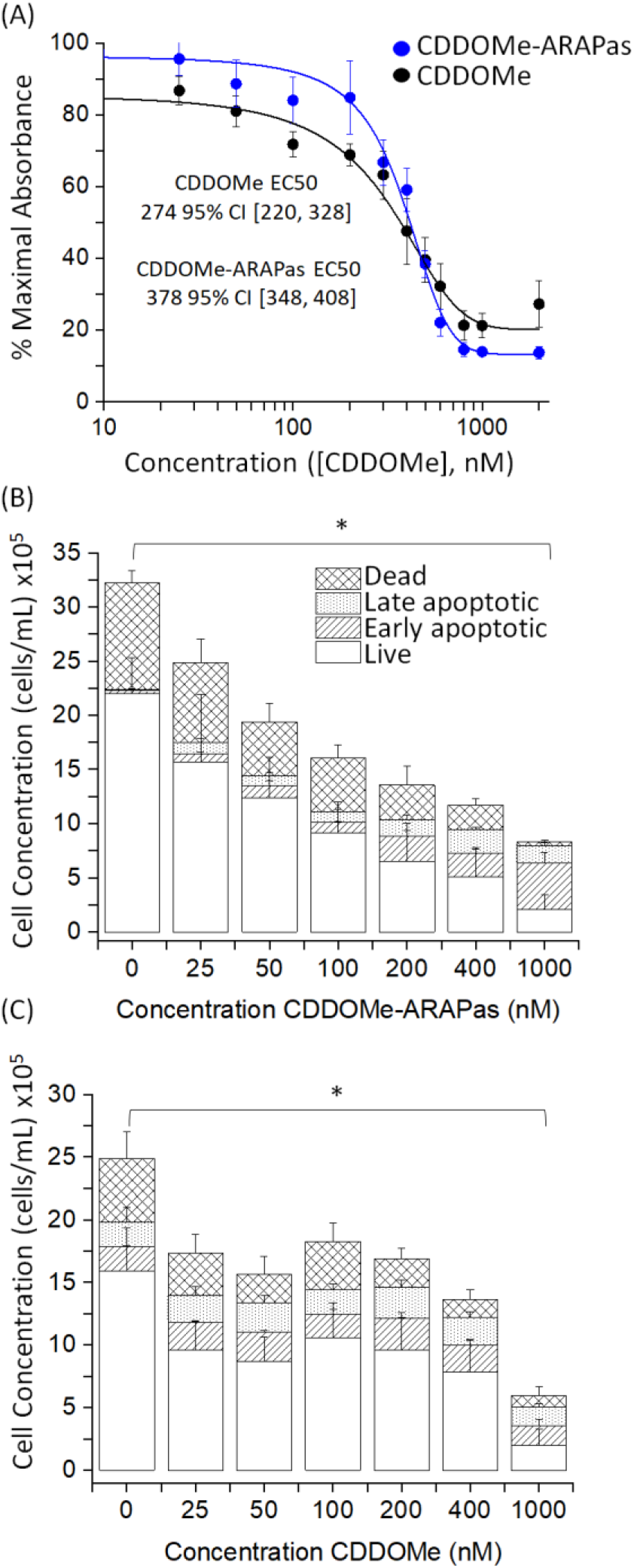
Toxicity of CDDOMe-ARAPas. **(A) MTT of RAW 264.7 macrophages treated with CDDOMe or CDDOMe-ARAPas**. Murine macrophages were incubated with CDDOMe or CDDOMe-ARAPas in a 0-2000nM dose range. Data represents N = 3-5 independent experiments ± SEM with 8 replicates per experiment. Data were fitted with a growth/sigmoidal curve with a dose response function in OriginPro 2018b software where the upper asymptote was fixed at 100 and the lower asymptote unfixed. The R square (COD) value for the CDDOMe-ARAPas and CDDOMe fitted curve were 0.98 and 0.99 respectively. The EC50 was derived from the fitted curves and were 274 95% CI [220, 328] and 378 95% CI [348, 408] for CDDOMe and CDDOMe-ARAPas respectively. **(B)-(C) Quantification of viable, early apoptotic, late apoptotic and dead RAW 264.7 macrophages following treatment with CDDOMe or CDDOMe-ARAPas**. Macrophages were incubated for 24 hours with 25-1000nM **(B) CDDOMe-ARAPas** or **(C) CDDOMe** and then cells were analyzed by the Muse Annexin V and Dead Cell Assay Kit. *p < 0.05 compared to control (0 µM CDDOMe-(ARAPas)). Data presented as means ± SEM (N = 3-5 independent experiments in triplicate). Data were analyzed using multivariate analysis of variance (MANOVA) to compare the effect of treatment with either CDDOMe or CDDOMe-ARAPas as well as concentration on total live cell population and then total dead cell population (where early and late apoptotic and dead cell populations were grouped together). Based on this analysis, we found a significant difference due to concentration, Wilk’s Lambda λ = 0.302, (F(12,62) = 4.24, P<0.001), with a large effect size estimate ω ^2^_Mult_ = 0.637. No difference was found due to treatment with either CDDOMe or CDDOMe-ARAPas. A separate univariate analysis (two-way ANOVA) revealed that the significant effect was upon the live cell population (including both CDDOMe and ARAPas) (F(6,32) = 5.72, P < 0.001), but not the dead cell population. A one-way ANOVA was conducted to examine the effect of concentration of either CDDOMe or CDDOMe-ARAPas on total live cell population. The one-way ANOVA for CDDOMe was significant (F(6, 14) = 3.21, P = 0.034) and for CDDOMe-ARAPas (F(6, 18) = 3, P = 0.033). Tukey post-hoc tests found a significant difference between the live cell population at 0 and 1000nM treatment in both CDDOMe (P = 0.04952) and CDDOMe-ARAPas (P = 0.02475).

### CDDOMe-ARAPas inhibit inflammation in classically activated macrophages

Previous studies by Kobayashi and co-workers reported that Nrf2 negatively regulates *iNOS* expression as well as that of pro-inflammatory cytokine genes, including *IL6* and *IL1b*; inhibiting LPS-induced expression of these genes in murine bone-marrow derived macrophages^20^. Moreover, synthetic triterpenoids such as CDDOMe were initially identified as potent inhibitors of iNOS protein expression in murine macrophages^42^. Thus, we sought to demonstrate that CDDOMe and, more importantly, the corresponding CDDOMe-ARAPas could negatively regulate iNOS protein expression. We also examined whether these could inhibit transcription of *IL1b* and *Il6*. Firstly, we examined *iNOS* mRNA levels with digital droplet PCR (ddPCR) in classically stimulated (IFNλ/LPS) murine RAW 264.7 macrophages in the presence or absence of CDDOMe or CDDOMe-ARAPas after 4 (**Fig. 5A**) or 24 hr incubation (**Supp. Fig. 1A**). A factorial ANOVA was conducted to determine the effect of either treatment with CDDOMe-ARAPas or CDDOMe alone, dose and stimulation with IFNλ/LPS on mRNA iNOS expression in macrophages. The model was significant (F(7, 19) = 67.59, P < 0.0001), and treatment (F(2, 19) = 76.62, P < 0.0001), dose (F(2, 19) = 43.73, P < 0.0001) and stimulation (F(1, 19) = 269.63, P < 0.001) were all significant. The effect size of treatment was ω^2^_p_ = 0.85, and for dose ω^2^_p_ = 0.76, whilst for IFNλ/LPS stimulation it was ω^2^_p_ = 0.91. Moreover, the interaction of dose and treatment with ARAPas vs CDDOMe alone was significant (F(2, 19) = 8.623, P = 0.002). Additionally, post hoc Tukey tests found that there were significant decreases in iNOS mRNA expression between vehicle and both 200 and 400nM of either CDDOMe alone or CDDOMe-ARAPas (**Fig. 5A**). At the 24 hr time point we found that overall iNOS mRNA expression trended to continue to be inhibited by treatment with CDDOMe at 400nM (**Supp. Fig. 1A**). However, there were no significant differences by factorial ANOVA as the overall model was not significant (F(5,12) = 2.33, P = 0.11). Secondly, we examined the production of nitric oxide by measuring nitrite in the media, as an index of iNOS protein activity (**Fig. 5B**). We conducted a factorial ANOVA analyzing the effects of macrophage stimulation with IFNλ/LPS, treatment with either CDDOMe-ARAPas or CDDOMe alone as well as dose upon picomol of NO / cell. Overall, this model was significant (F(7, 34) = 24.824, P < 0.0001), and analysis of the main effects showed that macrophage stimulation indeed significantly impacted NO/ cell (F(1, 34) = 76.04, P < 0.0001) with an effect size of ω^2^_p_ = 0.5. In line with this, a Tukey post-hoc test showed that for each vehicle treatment, classical stimulation conditions resulted in a production of picomol of NO / cell that was significantly increased (P < 0.0001), but that addition of vehicles themselves in the absence of IFNλ/LPS did not result in increased NO production. Furthermore, the main effects analysis showed that there was no statistically significant effect comparing treatment between CDDOMe vs CDDOMe-ARAPas (F(2, 34) = 3.05, P 0.061), however dose did have a statistically significant effect upon picomol of NO / cell produced (F(2, 34) = 21.084, P < 0.0001) with an effect size of ω^2^_p_ = 0.63. Moreover, a Tukey post-hoc test found a significant difference on picomol of NO / cell between the vehicle and 400nM for CDDOMe-ARAPas (P = 0.04664) and CDDOMe (P = 0.00105). Finally, we quantified iNOS protein levels via immunofluorescence (**Fig. 5C,D**). Classically stimulated murine RAW 264.7 macrophages were fixed, permeabilized and stained for iNOS protein expression, which was quantified as thresholded fluorescence per cell (RFU/cell). For each experiment, the RFU/cell for the IFNλ/LPS treated macrophages with the appropriate control vehicle (1% DMSO or polymer/PBS) was taken as the maximal RFU/cell, and relative RFU/cell for treatments (CDDOMe or CDDOMe-ARAPas) was calculated as a percentage of this value (**Fig. 5D**). We conducted a 2-way ANOVA to analyze the effect of treatment (with either CDDOMe-ARAPas or CDDOMe) or dose on percentage relative RFU/cell. The 2-way ANOVA model was significant (F(5, 10) = 20.61, P = 0.00006). Simple main effects analysis showed that there was no statistically significant effect comparing cells treated with CDDOme vs CDDOMe-ARAPas (F(1,10) = 3.92, P = 0.08), however dose did have a statistically significant effect upon relative RFU/cell (F(1, 10) = 33.38, P = 0.00004) with an effect size of ω^2^_p_ = 0.8. Additionally, the interaction between the two independent variables was significant (F(2, 10) = 4.93, P = 0.032) with an effect size of ω^2^_p_ = 0.33. Moreover, a Tukey post-hoc test found a significant difference on percentage relative RFU/Cell between the vehicle and 400nM for CDDOMe-ARAPas (P = 0.01236) and CDDOMe (P = 0.005). Our results are completely in line with previous results reported by Khoo and co-workers using an analog of CDDO-Me, CDDO-Imidazole^40^. These authors reported that co-treatment with 100nM CDDO-Im and LPS/IFNγ in RAW 264.7 macrophages significantly inhibited the induction of iNOS mRNA expression at 6 and 16 hr^40^. Moreover, it significantly decreased iNOS protein expression following 20hr co-incubation with LPS/IFNγ.

Additionally, we quantified *IL6* and *IL1b* mRNA expression in classically stimulated macrophages in the presence or absence of CDDOMe or CDDOMe-ARAPas, following either 4 or 24 hr incubation (**Supp. Fig. 1B-D**). Overall, we found that treatment with either CDDOMe or CDDOMe-ARAPas did not significantly affect either IL6 or Il1b mRNA expression. Kobayashi and colleagues found that these genes were negatively regulated by the Nrf2 transcription factor^20^. Dayalan Naidu and co-workers reported that another synthetic CDDO analog inhibited Il1-β mRNA expression in LPS-stimulated (1ng/mL) primary murine peritoneal macrophages at 4 hr^43^, with this dependent on the Cys 151 present in the Keap1. In another study, Zheng and co-workersreported that in LPS-stimulated (100ng/mL) RAW 264.7 macrophages, pre-treated for 1 hr with CDDO-Me (500nM), IL-1β protein production following 8 and 24hr incubation periods was inhibited, although no statistical tests on this data were reported^44^. *In vivo*, CDDO Me administration has reduced IL1-β protein in mesenteric adipose tissue of high fat diet fed mice^45^, and mRNA expression in of IL-1β, IL-6 and TNF-α in the liver and epididymal fat of diet-induced diabetic mice^46^. Together these studies support that pharmacological Nrf2 activation can decrease IL-1β and IL-6 expression, however reductions both *in vitro* and *in vivo* may be influenced by the stimulatory conditions employed, and cell/tissue type, which may explain the lack of inhibition in our study.

**Figure 5.**
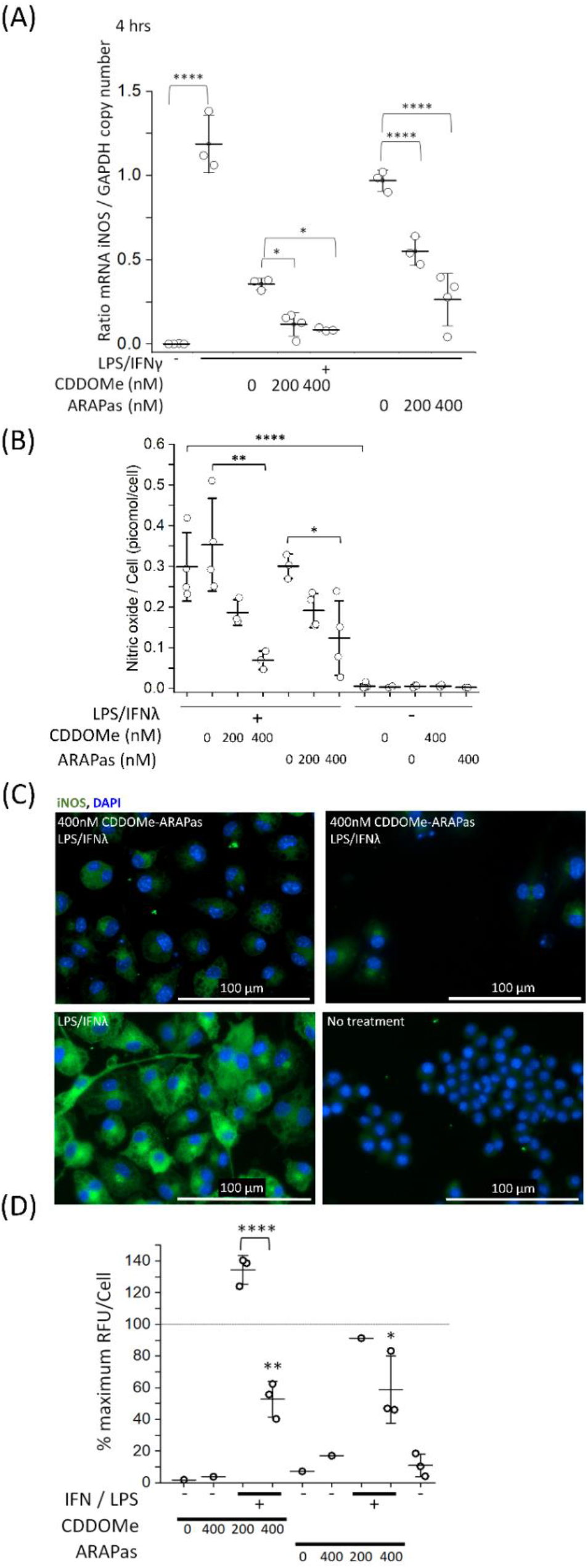
Inhibition of iNOS transcription and activity in classically stimulated macrophages. RAW 264.7 macrophages were classically stimulated with IFNλ (10ng/mL, 7 hr) followed by treatment with LPS (100 ng/mL, 4-24 hrs) in the presence or absence of treatments and their respective vehicles. The media was then collected and cells were either washed and scraped into sterile HBSS and pelleted for RNA extraction, or fixed (2% PFA), permeabilized (0.3% Triton X-100 in 10mM PBS) and then stained with anti-iNOS antibody followed by an appropriate secondary antibody. Secondary antibody alone and no antibody controls were included for all experiments. **(A) CDDOMe and CDDOMe-ARAPas dose-dependently inhibit iNOS mRNA expression after 4 hrs incubation**. Data represents the mean of N = 3-4 independent biological replicates, ± 1 S.D. A factorial ANOVA was conducted to determine the effect of either CDDOMe or CDDOMe-ARAPas treatment and dose upon iNOS mRNA expression normalized to GAPDH copy number (* P < 0.05, **** P < 0.0001). **(B) CDDOMe and CDDOme-ARAPas dose-dependently inhibit nitrite production after 24 hr incubation**. Nitrite in the media was reduced with acidic iodide reducing agent, and the resulting NO produced was detected with a Sievers NO analyzer. Cells were counted via DAPI staining, and NO normalized to cell count. Data represents the mean of 3-4 independent biological replicates, ± 1 S.D. A factorial ANOVA was conducted to determine the effect of either CDDOMe or CDDOMe-ARAPas treatment and dose upon picomol NO/cell (* P < 0.05, ** P < 0.01). **(C) Representative images of immunofluorescence of iNOS-stained murine macrophages after 24 hr incubation**. Fixed and permeabilized cells were stained with an anti-iNOS antibody followed by a secondary antibody conjugated to Alexa Fluor 488 dye, and then counter-stained with DAPI for nuclear detection. **(D) CDDOMe and CDDOMe-ARAPas inhibit iNOS protein expression in classically stimulated macrophages after 24 hr incubation**. Fluorescence intensity in the 488 channel was thresholded and normalized to cell count (RFU/Cell). For each treatment the respective vehicle+IFNλ/LPS was set as the maximal RFU/cell (100%), and percentage relative RFU/cell was determined for either CDDOMe or CDDOMeARAPas. Data represents the mean of N=1-3 biological replicates, with 3 biologcal replicates for all IFNλ/LPS treated conditions, conducted in sextuplicate, ± 1 S.D. A 2-way ANOVA was conducted to determine the effect of either CDDOMe or CDDOMe-ARAPas treatment and dose upon iNOS percentage relative RFU/Cell (* P < 0.05, ** P < 0.01, **** P < 0.0001).

### CDDOMe-ARAPas localize to atherosclerotic plaque

Athero-prone mice (LDLr^−/−^ and ApoE^−/−^) were high fat diet fed for 15 weeks in order to induce atherosclerotic plaque generation. To assess whether CDDOMe-ARAPas accumulated in atherosclerotic lesions, these mice were intravenously injected with 3 mg/kg of DiD-CDDOMe-ARAPas (25 μg/mL DiD dye). This was an equivalent dose of 125 μg/kg of DiD fluorescent dye. 24 hrs following injection these animals were sacrificed and the hearts, aortae, and a number of organs (liver, kidney, lung, gut, spleen) were excised for histology, and plasma collected. **Fig. 6A** shows representative fluorescence images of the aortic sinus region of ApoE^−/−^ and LDLr^−/−^ animals injected either with PBS or with 3 mg/kg of CDDOMe-ARAPas, whilst representative images of vehicle-injected animals are shown in **Supp. Fig. 2**. We quantified thresholded fluorescence intensity normalized to lesion area (RFU/lesion area, **Fig. 6B**) in the atherosclerotic plaque of the aortic sinus of both LDLr^−/−^ and ApoE^−/−^ mice. We conducted a 2-way ANOVA to determine the effect of both the genotype and treatment (DiD-CDDOMe-ARAPas vs vehicle) on RFU/lesion area. The 2-way ANOVA model was significant (F(3,11) = 7.41, P = 0.0055). Notably, we found that whilst genotype did not significantly affect RFU/lesion area ((F1,11) = 0.23, P = 0.64), treatment with CDDOMe-ARAPas did (F(1, 11) = 21.3, P = 0.0008), with an effect size of ω^2^_p_ = 0.57. Additionally the interaction between the variables was not significant (F(1, 11) = 0.04, P = 0.84). Overall, our data confirms that CDDOMe-ARAPas localize in atherosclerotic plaque.

**Figure 6.**
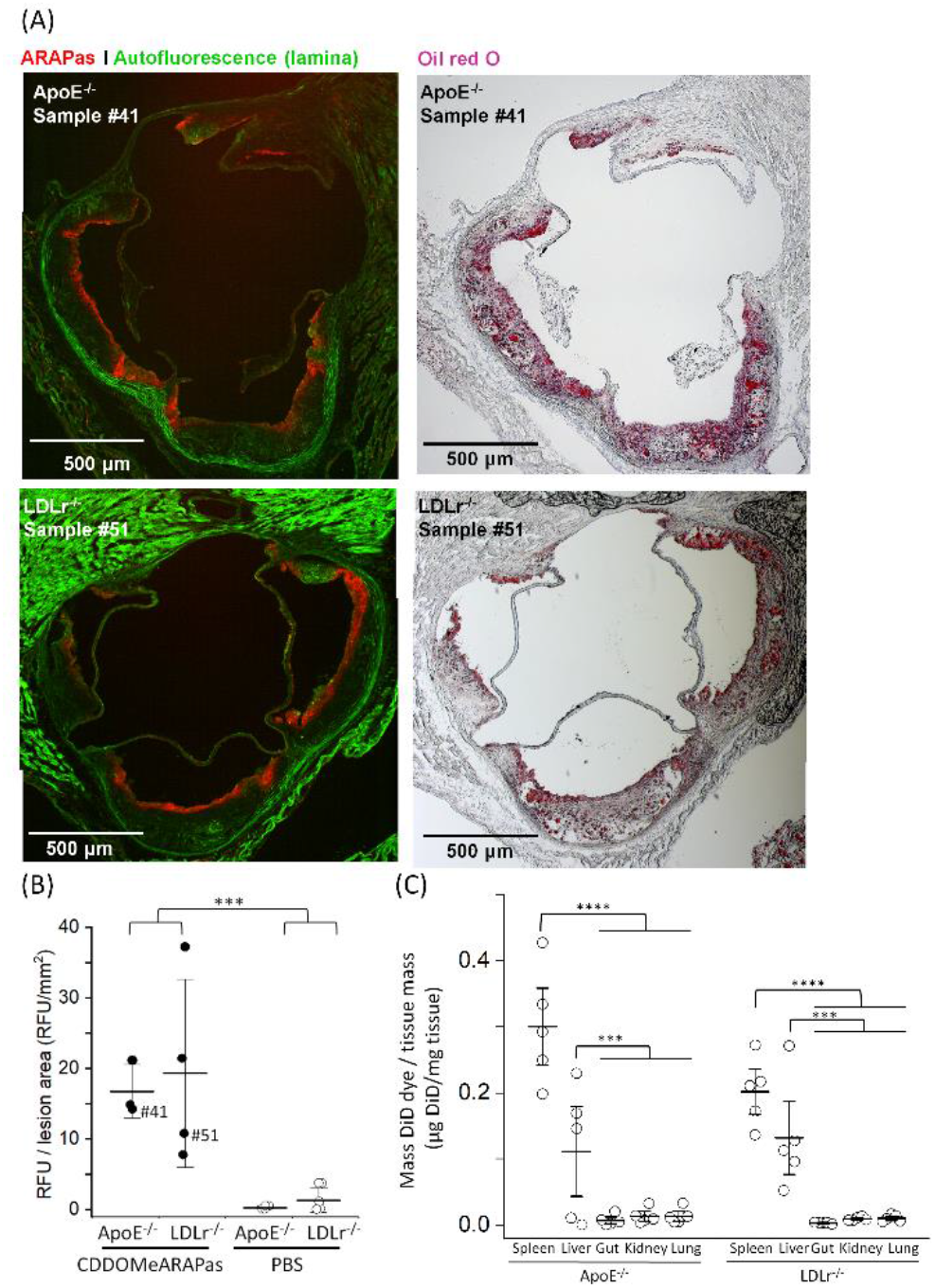
CDDOMe-ARAPas localize to atherosclerotic plaque in athero-prone mice. LDLr^−/−^ and ApoE^−/−^ mice were fed a high fat diet for 15 weeks and then intravenously injected with 3 mg/kg of DiD-CDDOMe-ARAPas (5mL/kg) or an equivalent volume of 10mM PBS. 24 hours later their organs and plasma were collected. **(A) Representative images of localization of DiD-CDDOMe-ARAPas in atherosclerotic plaque of both ApoE**^**-/-**^ **and LDLr**^**-/-**^ **mice**. Tissue sections were imaged by fluorescence microscopy and then the same tissue sectiosn were stained with Oil Red O to confirm the visualization of atherosclerotic plaque. **(B) CDDOMe-ARAPas selectively localize to atherosclerotic plaque. Aortic root sections were analyzed with fluorescence microscopy to determine localization of DiD-CDDOmeARAPas in atherosclerotic plaque**. DiD fluorescence intensity was thresholded and normalized to lesion area (determined by auto-fluorescence in the 488 channel). Data represents the mean of 3-5 independent biological replicates, ± 1 S.D. A factorial ANOVA was conducted to determine the effect of CDDOMe-ARAPas treatment and genotype upon RFU/lesion area (*** P < 0.001). **(C) Biodistribution of CDDOMe-ARAPas in athero-prone mice**. Organs were weighed and homogenized in 5% Triton-X-100 in 10mM PBS, and subsequently extracted with isopropanol. Fluorescence intensity was recorded and μg of DiD determined with a standard curve and normalized to tissue mass. Data represents the mean of 3-5 independent biological replicates, with fluorescence measurements conducted in triplicate, ± 1 S.D. Sham-injected genotype-matched animals were also measured and for each organ, the basal μg DiD/mg tissue was deducted. A factorial ANOVA was conducted to determine the effect of genotype and organs upon μg DiD/mg tissue (*** P < 0.001, **** P < 0.0001).

Nanoparticles take advantage of a dysfunctional endothelium with larger inter-endothelial junctions to accumulate in sites of vascular inflammation^14, 47^. Early atherosclerotic events have been shown to be associated with more severe endothelial barrier disruption, whilst advanced stage disease progressions are associated with improved endothelial barrier permeability^14^. Beldman and colleagues examined the junctional integrity of atherosclerotic lesions in both early (6 wk old) and advanced (12 wk old) plaque, and chiefly found that early atherosclerotic endothelial junctions could have spaces reaching up to 3μm in width, with slight but significant endothelial normalization for older more advanced lesions. Previous pre-clinical studies have successfully integrated nanoparticles into sites of atherosclerotic plaque, some of which were able to inhibit plaque formation and decreased the number of atherosclerotic lesions in mouse models of atherosclerosis^16, 47–51^. Macrophage-mediated uptake of nanoparticles *in vivo* is also likely to reflect phagocytosis due to both the presence of the protein corona that plays a role in nanoparticle-cell interactions^52^, and the nanoparticle size being >200 nm^15^. The accumulation of CDDOMe-ARAPas in our study is thus consistent with the literature, which shows that nanoparticles can accumulate selectively in atherosclerotic plaque due to the enhanced permeation and retention (EPR) effect^14, 47^. Overall, the mechanisms that influence nanoparticle uptake into atherosclerotic lesions is a continuing area of study. However, work by Beldman and co-workers^14, 53^ support a paradigm where intravenously injected nanoparticles chiefly enter atherosclerotic plaque through paracellular means by crossing the luminal endothelial junctions. Following paracellular transport, engulfment by plaque-associated macrophages is expected, which allows for nanoparticle retention in atherosclerotic plaque. We posit that this is also likely the means of plaque accumulation by the CDDO-Me-ARAPas.

In addition, we also examined their bio-distribution at 24 hr in each of the genotypes. To analyze their biodistribution, the organs were homogenized and the lysate extracted with isopropanol and DiD fluorescence measured with a fluorescence spectrophotometer. The average fluorescent signal from sham animals (injected with PBS vehicle) was subtracted from the signal in CDDOMe-ARAPas treated animals. We conducted a 2-way ANOVA to examine the influence of genotype and organs upon μg DiD /mg tissue. Overall, the 2-way ANOVA model was significant (F(9,40) = 19.5, P < 0.0001)), and it revealed that the effect of genotype was not significant (F(1,40) = 1.44, P = 0.24), whilst organs did influence μg DiD /mg tissue (F(4,40) = 41.6, P < 0.0001), with an effect size of ω^2^_p_ = 0.45. Moreover, post hoc Tukey tests revealed that μg DiD/mg tissue was significantly different particularly in the liver (P < 0.0001), and spleen (P < 0.001) compared to the other organs (kidney, lung, gut).

### Plasma chemistry profile *in vivo*

A number of plasma protein markers including aspartate transaminase (AST), alanine transaminase (ALT), and albumin (ALB) for liver toxicity, blood urea nitrogen (BUN) and creatinine for kidney toxicity, and finally cholesterol, low density lipoprotein, and triglycerides (TRIG) for a lipid profile, were measured. The data were analyzed using multivariate analysis of variance (MANOVA) to compare the effect of treatment with either CDDOMe-ARAPas or PBS and genotype (C57bl6, LDLr^−/−^ vs ApoE^−/−^) upon these markers (**Supp. Table 2**). The MANOVA revealed a significant difference due to treatment, Wilk’s Lambda λ = 0.337, (F(8,13) = 3.192, P = 0.031), and genotype, Wilk’s λ = 0.042 (F(16,26) = 6.26, P < 0.001), with a significant interaction between treatment and genotype, Wilk’s λ = 0.33 (F(8, 13) = 3.252, P = 0.029). The effect size of treatment upon ALT was estimate ω^2^_p_ = 0.53, and upon AST ω^2^_p_ = 0.67. Pairwise comparisons confirm that ApoE^−/−^ and LDLr^−/−^ have significantly increased levels of plasma cholesterol and LDL compared to those levels in age-matched normal chow-fed C57bl/6 mice, whilst triglyceride levels are only significantly increased in LDLr^−/−^ mice compared to both C57bl/6 (P = 0.001), and ApoE^-/^ (P < 0.0001). Regarding effects upon liver and kidney toxicity markers, pairwise comparisons revealed that CDDOMe-ARAPas treatment significantly increased ALT (P = 0.006) and AST (P < 0.0001) in LDLr^−/−^ mice, but without significant changes observed in ApoE^−/−^ mice. In ApoE^−/−^ mice, pairwise comparisons show a significant decrease in BUN (P = 0.028) and increase in creatinine (P = 0.026). All other markers did not show significant changes between treatments, or within genotypes. An increase in serum levels of AST and ALT was also observed in patients treated with CDDO-Me^54^. In this study the increase in serum levels of transaminases was transient. Even though the increase in serum levels of these enzymes is associated with hepatotoxic effects, Lewis and colleagues suggested that Nrf2 activation increases serum levels of AST and ALT by increasing their gene expression in both hepatic and extra-hepatic tissues^54^. The authors showed that serum levels of AST and ALT are lower in Nrf2-null mice and higher in Keap1-knockdown mice. Since the CDDO-Me-ARAPas accumulated significantly in the liver, it is possible that hepatic Nrf2 activation could drive AST and ALT gene expression. It would be important to assess Nrf2 activation status in the liver and assess whether the observed increased in AST and ALT is transient.

### CDDOMe-ARAPas locally activate Nrf2 in atherosclerotic plaque

Finally, we examined whether CDDOMe and CDDOMe-ARAPas activate Nrf2-regulated genes, GCLC and NQO1, in both classically stimulated murine macrophages (**Fig. 7A, B**) and in high fat diet fed LDLr^−/−^ mice (**Fig. 7C, D**). In classically stimulated murine macrophages incubated with CDDOMe or CDDOMe-ARAPas (200-400 nM) we found that GCLC (**Fig. 7A**) and NQO1 (**Fig. 7B**) mRNA expression increased. A factorial ANOVA was performed to analyze the effect of IFNλ/LPS stimulation, and CDDOMe or ARAPas treatment on GCLC mRNA expression. The model was significant (F(4,17)=36.4 P<0.001) and revealed that GCLC was decreased with classical stimulation (F(1,17)=143.6, P<0.001) with an effect size of ω^2^_p_ = 0.87. This is in line with literature reports that LPS stimulation decreases GCLC mRNA expression^55^. Simple main effects analysis revealed that treatment with either CDDOMe or CDDOMe-ARAPas had a significant effect on GCLC mRNA expression (F(2,17)=4.6 P=0.026) with an effect size of ω^2^_p_ = 0.14. However, the difference in concentration between 200 to 400nM was non-significant (F(1,17)=1.7 P=0.206). Overall, GCLC mRNA expression was increased by CDDOMe (1.7-1.9 fold) and by CDDOMe-ARAPas (3-4.5 fold) (**Fig. 7A**). Post hoc Tukey pairwise comparisons showed that these changes were statistically significant compared to control with P=0.031, and P=0.029, for both CDDOMe and CDDOME ARAPas, respectively (**Fig. 7A**). Regarding NQO1 mRNA expression in classically stimulated macrophages, we conducted a factorial ANOVA on whether treatment with CDDOMe or CDDOMe-ARAPas, and dose influenced NQO1 mRNA expression. This model was also significant (F(5,12) = 9.2, P < 0.0001), and revealed that both treatment (CDDOMe vs CDDOMe-ARAPas) (F(1,12) = 22.13, P < 0.0001) and dose was significant (F(2,12) = 14, P = 0.0014), with effect sizes of ω^2^_p_ = 0.54 and ω^2^_p_ = 0.55, respectively. Moreover, the interaction of these two variables was non-significant (F(2,12) = 3.2, P = 0.08). Post hoc Tukey pairwise comparisons showed that NQO1 mRNA expression was significantly increased by CDDOMe-ARAPas (2.7-fold) (**Fig. 7B**).

**Figure 7.**
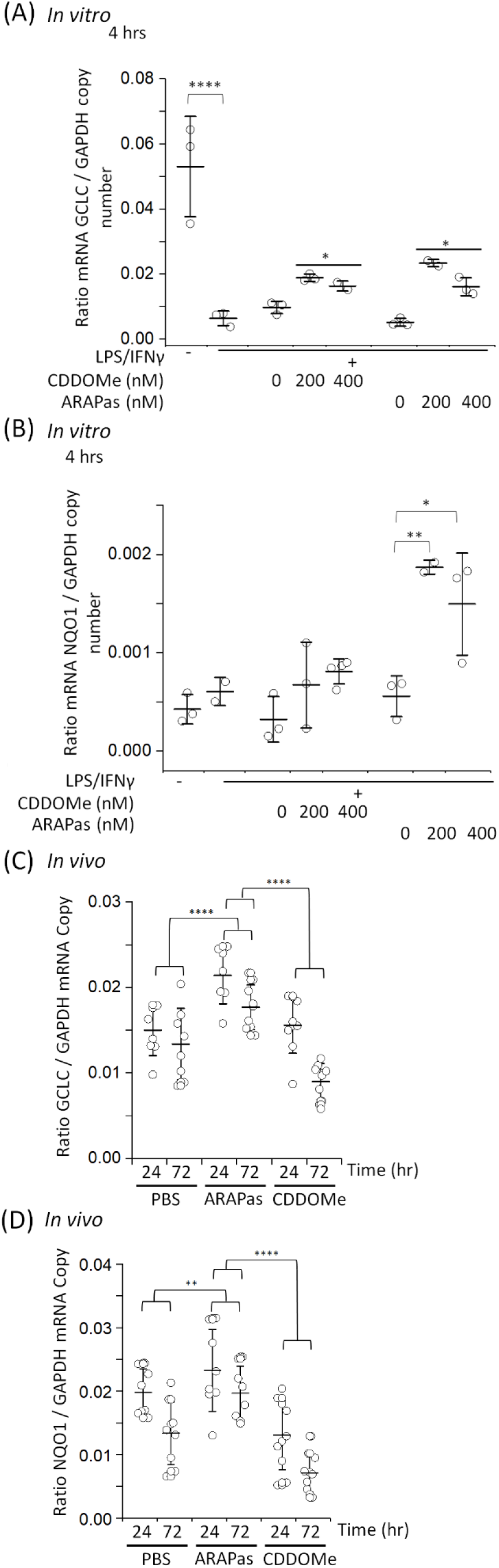
Activation of Nrf2-regulated genes *in vitro* and *in vivo* by CDDOMe-ARAPas. GCLC and NQO1 mRNA expression was recorded in either classically stimulated murine RAW 264.7 macrophages or in arotic arch homogenates of high fat diet fed LDLr^−/−^ mice. RAW 264.7 macrophages were classically stimulated with IFNλ (10 ng/mL, 7 hr) followed by treatment with LPS (100 ng/mL, 4 hrs) in the presence or absence of treatments and their respective vehicles. Cells were either washed and scraped into sterile HBSS and pelleted for RNA extraction. 4-6 wk old LDLr^−/−^ mice were high fat diet fed for 8 weeks (till 12-14 wks old) and then either injected intravenously with 1mg/kg of CDDOMe encapsulated in ARAPas or 10mM PBS, or intraperitoneally with 1mg/kg CDDOme in DMSO. Aortic arches were collected at either 24 hr or 72 hr post injection of CDDOMe-ARAPas, 10 mM PBS, or CDDOme. **(A) Activation of GCLC mRNA expression in classically activated murine macrophages following 4 hr incubation**. Data represents the mean of 2-3 independent biological replicates, ± 1 S.D. A factorial ANOVA was conducted to determine the effect of classical stimulation, treatment and dose upon GCLC mRNA expression normalized to GAPDH copy number (* P < 0.05, **** P < 0.0001). 200-400nM of either CDDOme or CDDOMe-ARAPas were significant compared to their respective vehicles. **(C) Activation of NQO1 mRNA expression in classically activated murine macrophages following 4 hr incubation**. Data represents the mean of 3-4 independent biological replicates, ± 1 S.D. A factorial ANOVA was conducted to determine the effect of treatment with either CDDOme or CDDOMe-ARAPas and dose upon NQO1 mRNA expression normalized to GAPDH copy number (* P < 0.05, ** P < 0.01). **(C) Activation of GCLC mRNA expression in LDLr**^**-/-**^ **aortic arch homegenates**. Data represents the mean of 8-10 independent biological replicates for each condition, ± 1 S.D. A factorial ANOVA was conducted to determine the effect of treatment with either PBS, CDDOme or CDDOMe-ARAPas and time upon GCLC mRNA expression normalized to GAPDH copy number (**** P < 0.0001). **(C) Activation of GCLC mRNA expression in LDLr**^**-/-**^ **aortic arch homegenates**. Data represents the mean of 8-10 independent biological replicates for each condition, ± 1 S.D. A factorial ANOVA was conducted to determine the effect of treatment with either PBS, CDDOme or CDDOMe-ARAPas and time upon NQO1 mRNA expression normalized to GAPDH copy number (** P < 0.01, **** P < 0.0001).

We next wanted to confirm whether selective delivery of CDDOMe-ARAPas to atherosclerotic plaque (as shown in **Fig. 6**) led to selective increases in mRNA expression of Nrf2-regulated genes GCLC and NQO1. LDLr^−/−^ mice were high fat diet fed over 8 weeks and then intravenously injected with either 10mM PBS, or 1mg/kg CDDOMe encapsulated in ARAPas (equivalent to ~3mg/kg nanoparticle mass). HPLC-UV-VIS was used to determine the concentration of CDDOMe in ARAPas suspensions, and the volume injected was adjusted to ensure a dose of 1mg/kg. To compare to un-targeted CDDOMe, we injected an equi-molar dose of CDDOMe (1mg/kg) intra-peritoneally, dissolved in DMSO. Typically, the doses cited in the literature for CDDOMe treatment range from 3-100 mg/kg^56–58^, with a majority employing 10mg/kg. We chose to use a lower dose than previously reported to better parse out whether encapsulation of CDDOMe in ARAPas, resulting in localized delivery to atherosclerotic plaque, would result in localized Nrf2 activation that is absent in un-targeted CDDOMe. 24 and 72 hours following injection, the aortic arch (a site of preferential atherosclerotic plaque formation in mice) was harvested and subsequently homogenized and mRNA extracted and purified. GCLC (**Fig. 7C**) and NQO1 (**Fig. 7D**) mRNA in aortic arch homogenates were detected utilizing ddPCR. IFNλ/LPS treatment significantly decreased GCLC mRNA expression as previously published^59^. Firstly, we conducted a 2-way ANOVA to determine whether treatment (PBS, CDDOMe-ARAPas, CDDOMe alone) and time (24 vs 72 hrs) influenced GCLC mRNA expression normalized to GAPDH copy number. The model was significant (F(5,45) = 15.5, P < 0.0001), and simple main effects analysis revealed that both treatment (F(2,45) = 25, P < 0.0001) and time (F(1,45) = 21, P < 0.0001) significantly influenced GCLC mRNA expression, although the interaction was non-significant (F(2,45) = 3, P = 0.07). Overall, the effect size for treatment was ω^2^_p_ = 0.48, and for time ω^2^_p_ = 0.28. Post-hoc Tukey tests showed that there was a significant increase in GCLC mRNA expression for CDDOMe-ARAPas vs PBS (P < 0.0001) and for CDDOMe-ARAPas vs CDDOMe alone (P < 0.0001) (**Fig. 7C**). On the other hand, there was a non-significant change in GCLC expression for CDDOMe alone vs PBS (P = 0.13). To examine the changes in NQO1, we also conducted a 2-way ANOVA with treatment (PBS, CDDOMe-ARAPas, CDDOMe alone) and time (24 vs 72 hrs). The model was significant (F(5,47) = 14, P < 0.0001), and simple main effects analysis revealed that both treatment (F(1,47) = 16.5, P < 0.0001) and time (F(2,47) = 26.2, P < 0.0001) significantly influenced NQO1 mRNA expression, although the interaction was non-significant (F(2,47) = 0.42, P = 0.66). Overall, the effect size for treatment was ω^2^_p_ = 0.37, and for time ω^2^_p_ = 0.32. Post-hoc Tukey tests showed that there was an increase in mRNA expression for CDDOMe-ARAPas vs PBS (P < 0.00969) and for CDDOMe-ARAPas vs CDDOMe alone (P < 0.0001) (**Fig. 7D**). For NQO1, there was also a significant difference in mRNA expression for CDDOMe alone vs PBS (P < 0.0001), with CDDOMe alone decreasing expression. Khoo and co-workers previously found that an analog of CDDO-Me, CDDO-imidazole, was able to dose-dependently increase HO1 expression in naïve RAW 264.7 macrophages, as well as significantly increase HO1 expression above that of the HO1 already expressed in IFNλ/LPS treated RAW 264.7 macrophages^40^. Thus, we also investigated the effect of CDDOMe and CDDOMe-ARAPas injection on HO1 and SOD1 mRNA expression at 24 and 72 hours post-injection (**Supp. Fig. 3**), however we did not find a change to the expression of these mRNA for any treatment *in vivo*.

In the literature, several compounds known as Nrf2-inducers (tBHQ^22^, Ebselen^23^, CDDO-Me analogue dh404^24, 56^, and oleanolic acid^25^) have augmented endogenous antioxidant systems and limited inflammation thereby preventing atherosclerosis development or progression in diabetes-aggravated atherosclerosis^56^. In particular, Tan and colleagues using a related synthetic triterpenoid analog, dh404, examined mRNA levels of NQO1, Glutathione S Transferase (GSH-S-T) and Glutathione peroxidase 1 (GPx1) in the kidney cortex following 5 weeks of treatment. In this study, only the 20mg/kg dose (not 3 or 10) had increased GSH-S-T, and GPx1 mRNA in diabetic animal kidney cortex relative to the vehicle^56^. Overall, the results presented in **Fig. 7C** and **D** support that selective delivery of CDDOMe-ARAPas to atherosclerotic plaque results in a targeted increase in mRNA expression of Nrf2-regulated genes in the aortic arch, which is absent in equi-molar untargeted CDDOMe. In light of the reported anti-inflammatory and athero-protective nature of pharmacological Nrf2 induction, we speculate that localized delivery will allow for greater targeted effect with lower doses. A barrier for the clinical use of redox-modulatory drugs is their adequate access to sites of redox dysfunction such as atherosclerotic plaque, thus our study begins to address this important challenge.

## Conclusions

Herein we report the successful encapsulation of the potent Nrf2 activator drug, CDDO-Methyl into polymeric nanoparticles via flash nanoprecipitation (FNP)^29^ to generate CDDOMe Antioxidant Response Activating nanoParticles (CDDOMe-ARAPas). FNP was chosen as our nanoparticle synthesis method as it is highly translational, being amenable to large-scale (kg/day) manufacturing whilst robustly maintaining nanoparticle characteristics and homogeneity^30^. We describe the physiochemical characteristics of our nanoparticles as well as their *in vitro* release of CDDO-Me, resulting in potent Nrf2 activation. We additionally demonstrate their internalization by naïve murine macrophages and inhibition of pro-inflammatory iNOS induction by classically activated macrophages. We go on to show that these nanoparticles selectively accumulate in atherosclerotic plaque in two widely used genotypes of athero-prone mice (ApoE^−/−^ and LDLr^−/−^). Finally, we show that the CDDOMe-ARAPas successfully activate the expression of Nrf2-regulated genes both *in vitro* in murine macrophages and *in vivo* in the aortic arch. Moreover, equi-molar doses of un-encapsulated CDDOMe fail to induce expression under the same conditions *in vivo*. It was beyond the scope of this study to assess whether intra-plaque delivered Nrf2 activator drugs such as CDDO-Me, prevent atherosclerotic progression, which will be assessed in future work. Overall, these studies demonstrate the successful intra-plaque delivery of antioxidant-based therapeutics employing a highly translational nanoparticle synthesis technique. Our studies support the paradigm that targeted delivery of redox-active therapeutics is superior to systemic delivery for modulation of the intra-plaque environment^20^.

## Supporting information

Electronic Supplementary Information

## Author Contributions

**Sophie Maiocchi:** Conceptualization, Methodology, Formal analysis, Investigation, Data curation, Writing – original draft, Visualization. **Ana Cartaya:** Investigation, Visualization. **Sydney Thai:** Investigation, Formal analysis, Data curation. **Adam Akerman:** Methodology, Resource. **Edward Bahnson:** Conceptualization, methodology, Formal Analysis, Resources, Writing – Review & Editing, Supervision, Project Administration, Funding Acquisition.

## Acknowledgements

This work was supported by the National Institutes of Health, National Heart Lung and Blood Institute [K01HL145354] to E.M.B. S. M. is supported by the Leon and Bertha Golberg Fellowship. A. E. C. is supported by National Heart Lung and Blood Institute [F31HL156427] We gratefully acknowledge Vicky Madden and Kristen White for assistance with conventional TEM. The Microscopy Services Laboratory, Department of Pathology and Laboratory Medicine, is supported in part by P30 CA016086 Cancer Center Core Support Grant to the UNC Lineberger Comprehensive Cancer Center. We also gratefully acknowledge Marina Sokolsky, Olesia Gololobova and Jacob Ramsey of the Nanomedicines Characterization Core Facility (NCore) at the Center for Nanotechnology in Drug Delivery (CNDD) at UNC School of Pharmacy for their assistance with Nanosight Nanotracking Analysis and HPLC UV-VIS analysis of nanoparticle samples. Additionally, we thank Ling Wang in the Animal Histopathology and Laboratory Medicine Core for expert technical assistance with analysis of plasma chemistry. The Animal Histopathology and Laboratory Medicine core is supported in part by an NCI Center Core Support Grant (5P30CA016080-42). Confocal imaging was performed at the UNC Neuroscience Microscopy Core supported, in part, by the NIH-NINDS P30 NS045892 and the NIH-NICHD U54 HD079124.

