## Supplementary material for "Antioxidant Response Activating nanoParticles (ARAPas) localize to atherosclerotic plaque and locally activate the Nrf2 pathway": Electronic Supplementary Information

Received 00th January 20xx,
Accepted 00th January 20xx

DOI: 10.1039/x0xx00000x

**Sophie Maiocchi,[a,b,c,d] Ana Cartaya,[c,d,e] Sydney Thai,[f] Adam Akerman[f] and Edward Bahnson[a-e,*]**

Atherosclerotic disease is the leading cause of death world-wide with few novel therapies available despite the ongoing health burden. Redox dysfunction is a well-established driver of atherosclerotic progression; however, the clinical translation of redox-based therapies is lacking. One of the challenges facing redox-based therapies is their targeted delivery to cellular domains of redox dysregulation. In the current study, we sought to develop Antioxidant Response Activating nanoParticles (ARAPas), encapsulating redox-based interventions, that exploit macrophage biology and the dysfunctional endothelium in order to selectively accumulate in atherosclerotic plaque. We employed flash nanoprecipitation (FNP) to synthesize bio-compatible polymeric nanoparticles encapsulating the hydrophobic Nrf2 activator drug, CDDO-Methyl (CDDOMe-ARAPas). Nuclear factor erythroid 2-related factor 2 (Nrf2)-activators are a promising class of redox-active drug molecules whereby activation of Nrf2 results in the expression of several antioxidant and cyto-protective enzymes that can be athero-protective. In this study, we characterize the physiochemical properties of CDDOMe-ARAPas as well as confirm their in vitro internalization by murine macrophages. Drug release of CDDOMe was determined by Nrf2-driven GFP fluorescence. Moreover, we show that these CDDOMe-ARAPas exert anti-inflammatory effects in classically activated macrophages. Finally, we show that CDDOMe-ARAPas selectively accumulate in atherosclerotic plaque of two widely-used murine models of atherosclerosis: $ApoE^{-/-}$ and $LDLr^{-/-}$ mice, and are capable of increasing gene expression of Nrf2-transcriptional targets in the atherosclerotic aortic arch. Future work will assess the therapeutic efficacy of intra-plaque Nrf2 activation with CDDOMe-ARAPas to inhibit atherosclerotic plaque progression. Overall, our present studies underline that targeting of atherosclerotic plaque is an effective means to enhance delivery of redox-based interventions.

#### Extended Experimental

##### TEM

Conventional TEM images were captured on a JEOL JEM 1230 TEM at 80 kV, LaB6 filament, with a Gatan Orius SC1000 CCD camera and Gatan Microscopy Suite 3.0 software. CDDOMe-ARAPas at 0.3 mg/mL in phosphate buffered saline (PBS) were prepared for TEM by pipetting 8 µL atop Formvar on copper 400 mesh TEM grids (Ted Pella) that were treated with glow discharge (PELCO easiGlowTM Glow Discharge Cleaning System). After two minutes, samples were rinsed with deionized water and stained with 2% uranyl acetate negative stain for two minutes prior to imaging.

##### Nanosight nanotracking

The nanoparticle hydrodynamic diameter size via particles/mL distribution was measured using the NanoSight NS500 (Malvern Panalytical, Ltd, UK) and Zetaview QUATT – NTA Nanoparticle Tracking – Video Microscope PMX-420 (Particle Metrix). Samples were prepared in either 5mM Citrate Buffer (pH 5) or 10mM PBS (pH 7.4) at 0.01-0.005mg/mL. All buffers were 0.02µm filtered prior to use. At least 5 measurements of 60 s were taken per sample at 25°C.

##### DLS

The nanoparticle hydrodynamic diameter size via intensity distribution and polydispersity index were characterized using a Zetasizer Nano ZS (Malvern Panalytical, Ltd, UK). Samples were prepared in 10mM PBS at 0.05mg/mL. At least 10 scans were taken per measurement, three measurements per sample at 25°C.

##### HPLC-UV-VIS

CDDO-Me loading capacity and final concentration in nanoparticle suspensions were obtained using an Agilent High Performance Liquid Chromatography (HPLC) system with a UV-VIS detector (Agilent Technology 1200 series). CDDO-Me was resolved with a Supelco Analytical Nucleosil C18 HPLC column (Cat# Z226181, 25 cm x 4.6 mm, 5µm particle size). The area under the peak was recorded for the 200 nm wavelength filter. The mobile phase consisted of A: 0. 1% (v/v) trifluoroacetic acid in Acetonitrile and B: 0. 1% (v/v) trifluoroacetic acid in water, at an isocratic flow of 1 mL/min consisting of 85% A. All mobile phase solvents were HPLC-grade. Samples, 10 µl, were prepared

[a.] *Department of Cell Biology and Physiology. University of North Carolina at Chapel Hill, NC 27599.*
[b.] *Curriculum of Toxicology and Environmental Medicine. University of North Carolina at Chapel Hill, NC 27599*
[c.] *Center for Nanotechnology in Drug Delivery. University of North Carolina at Chapel Hill, NC 27599*
[d.] *McAllister Heart Institute. University of North Carolina at Chapel Hill, NC 27599.*
[e.] *Department of Pharmacology. University of North Carolina at Chapel Hill, NC 27599.*
[f.] *Department of Surgery. University of North Carolina at Chapel Hill, NC 27599.*

Please do not adjust margins



as 5-50µg/mL solutions in 90% HPLC-grade acetonitrile. A standard curve with CDDO-Me was conducted each time.

**Macrophage Internalization Assay**

RAW 264.7 macrophages were seeded at 500,000 cells per well in a 12-well corning sterile cell culture plate. Cells were incubated overnight in high glucose DMEM (Gibco) supplemented with 10% FBS, 1% penicillin/streptomycin). Cells were washed with HBSS and then treated with fluorescent DiD-loaded CDDOMe-ARAPas to a final concentration of 5µg/mL DiD in media or equivalent amounts of PBS. Cells were incubated at either 4°C or 37°C for 18 hr. At the end of the incubation period, cells were washed with cold HBSS and then lysed with cold 1% Triton-X-100 in PBS (1mL per well). Cells were scraped, centrifuged (14,000 rpm, 15 min, 4°C). Cell lysate was measured for its fluorescence (excitation $\lambda$ = 640nm, emission $\lambda$ = 690nm) using quartz cuvettes (Hellma fluorescence cuvette, Micro-suprasil, Cat# Z802875) and a spectrofluorometer (Spectra Max M3) and recorded in SoftMax Pro Software. Fluorescence was converted to µg/mL DiD with a standard curve. Cell lysate was measured for protein in µg/mL using the BCA assay (thermo fisher). Results were normalized and expressed as µg of DiD / µg of protein.

**Macrophage Association Assay**

RAW 264.7 macrophages were seeded at 50,000 cells per well in a 96 well black glass-bottom plate (Greiner 96 well plates, 655891, VWR). Cells were treated with 10µg/mL Hoescht dye for 30 minutes at 37°C, 5% CO2. Cells were then imaged every 4 hours continuously over 18 hours after having media replaced with DMEM no phenol red media (10% FBS, 1% penicillin/streptomycin) containing treatments (PBS (20%) or CDDOMe-ARAPas; final concentration of 0.1mg/mL). Wells were imaged using Gen 5 software (BioTek Instruments, Winooski, VT) on a Cytation 5 plate reader (BioTek Instruments) (37°C, 5% CO2) with a DAPI filter cube (BioTek Instruments, part #:1225100) and a Texas Red filter cube (BioTek Instruments: Part # 1225102). Exposure times were fixed for each well. Cells were counted automatically by the Gen 5 software by thresholding in the DAPI channel.

**Classical Activation of Macrophages**

Murine Raw 264.7 macrophages were plated at 20,000 cells/well in a 96-well glass bottom plate (Greiner Cellview microplate, Cat#: 655891), and allowed to attach overnight in complete media (DMEM High glucose, Gibco, 1% penillin/streptomycin, 10% Heat inactivated FBS). Cells were primed with IFNλ (10ng/mL, R&D Systems, 485-MI) for 7 hrs, prior to addition of treatments with LPS (100ng/mL, Sigma-aldrich, L4391-1MG)[32].

**MTT assay**

RAW 264.7 macrophages were seeded in 96-well plates (5000/ well). Macrophages were treated with a range of concentrations (0-2000 nM) of either CDDOMe or CDDOMe-ARAPas for 24 hrs. RAW macrophages were then treated with 0.4 mg/mL Thiazolyl Blue tetrazolium bromide (L11939; Alfa Aesar, Haverhill, MA) diluted in high glucose FBS and Pen-Strep-supplemented DMEM for 4 h at 37 °C. After treatment, media was removed and plates were left to air-dry overnight. Formazan crystals were resuspended in DMSO and absorbance was measured at 560 nm with background at 670 nm on a Cytation 5 plate reader.

**Cell apoptosis assay**

RAW 264.7 macrophages were seeded in 12-well plates (1 x $10^5$ / well). These were treated with 25-1000 nM CDDOMe or CDDOMe-ARAPas for 24 h. Cells were scraped into sterile cold HBSS, pelleted by centrifugation (300×*g*, 5 min), and stained using the manufacturer's instructions (MCH100105, Sigma-Aldrich). Live, early apoptotic, late apoptotic, and dead cells were quantified for each well using Muse Cell Analyzer. Each proportion was determined by the appropriate combination of negative or positive staining for Annexin V and 7-Aminoactinomycin D.

**Measurement of nitrite in complete media samples.**

Cells were treated for 24 hr, at which point the media was removed for analysis of nitrite levels. Cells were washed with ice-cold HBSS, and then fixed with 2% PFA in PBS for 15 min. Cells were stained with DAPI (1/500 dilution) for 5 min and then imaged with a Cytation 5.0 Plate Reader to obtain the total cell count per well. Nitrite in complete media samples was analyzed by iodide chemiluminescence with the Sievers 280i Nitric Oxide Analyzer (NOA) and the corresponding packaged software Liquid.vi software as per the NO Analyzer manufacturer instructions. Acidic iodide reducing reagent was prepared fresh (50mg potassium iodide, 2mL distilled H2O and 5mL glacial acetic acid). A standard curve was conducted with sodium nitrite solutions prior to sample measurements. Complete media samples were diluted with ice-cold PBS (1 volume of complete media to 1 volume of PBS). 50µL of diluted complete media samples was injected into the reducing reagent and the area under the peak was used to measure the content of nitrite in picomol.

**Measurement of iNOS with immunofluorescence**

Polarized RAW 254.7 macrophages, treated for 24 hr, were washed with ice-cold HBSS, and then fixed with 2% PFA in PBS for 15 min, and permeabilized with 0.3% Triton-X-100 in 1XPBS for 15 min. Cells were stained with a rabbit polyclonal anti-iNOS antibody (ab3523), and then an AF488 goat anti-rabbit secondary antibody, and DAPI (1/500 dilution) and then imaged using Gen 5 software (BioTek Instruments, Winooski, VT) on a Cytation 5 plate reader (BioTek Instruments) with a DAPI filter cube (BioTek Instruments, part #:1225100) and a GFP filter cube (BioTek Instruments, part #:1225101). Exposure times were fixed for each well. Cells were counted automatically by the Gen 5 software by thresholding in the DAPI channel.

 This journal is © The Royal Society of Chemistry 20xx

Please do not adjust margins

Please do not adjust margins



**Measurement of gene expression with Digital Droplet PCR (ddPCR)**

**RNA Copy Number Quantification**

Total RNA was extracted by ethanol precipitation using TRIzol Reagent (15596026, Thermo Fisher Scientific) and quantified by the Cytation 5.0 plate reader with the Take3 microvolume plate (BioTek). For mRNA, cDNA was synthesized from 1ug of total RNA using the qScript cDNA synthesis kit (95047, Quantabio); prior to PCR amplification of mRNAs, cDNA was diluted in RNase-free water (1:10 – 1:30). These dilutions allowed for a minimum count of 100 copies of each RNA copy to be measured in each ddPCR reaction. For each ddPCR assay, 5 uL cDNA sample, 12.5uL ddPCR supermix for probes (no dUTP) (Bio-Rad, 1863024), and 5 uL RNase-free Water was added to the mixture. For murine mRNAs, a 1X concentration of the following FAM labeled probes (HO1, NQO1, GCLC, SOD1, iNOS, IL6, and IL1b) were multiplexed with VIC labeled Glyceraldehyde 3-phosphate dehydrogenase (GAPDH) probe. All primer and gene target information is available in **Supplementary Table 1**. Information concerning specificity, linear dynamic range, sensitivity, and efficiency in gene expression analyses is available online at: https://www.thermofisher.com/order/genome-database/ and can be found for each TaqMan Gene Expression Assay by searching the catalogue numbers provided in Supplementary Table 1. The mixture and 70 uL of droplet generation oil for probes (Bio-Rad, 1863005) were loaded into the sample and oil wells of a disposable droplet generator cartridge (1864008, Bio-Rad). Droplets were generated by the QX-200 droplet generator (Bio-Rad) and carefully transferred to a 96-well PCR plate by pipetting the mixture down the side of the well at an angle. The plate was sealed with foil at 180 °C. The thermo cycling conditions were: 95°C for 10 minutes, 94°C for 30 seconds, 60°C for 1 minute, repeat 39 times from step 2, 98°C for 10 minutes, 4°C for 30 minutes. After which the plate was incubated at room temperature for 5 minutes. Droplets were read in the QX200 droplet reader (Bio-Rad). Only samples containing greater than 10,000 droplets were included in analysis of copy number determination. Poisson distribution was used to determine the number of template molecules per droplet using Quantasoft Analysis Pro (Bio-Rad). Normalization was performed by dividing all copy numbers by the internal controls GAPDH values.

**Supplementary Table 1.**

| **Species** | **Gene Name** | **Gene Symbol** | **Assay ID (Thermo Fisher Scientific)** |
|---|---|---|---|
| mouse | NAD(P)H dehydrogenase, quinone 1 | *nqo1* | Mm01253561_m1 |
| mouse | superoxide dismutase 1 | *sod1* | Mm01700393_g1 |
| mouse | glutamate-cysteine ligase, catalytic subunit | *gclc* | Mm00802655_m1 |
| mouse | heme oxygenase 1 | *hmox1* | Mm00516005_m1 |
| mouse | interleukin 1 beta | *il1b* | Mm00434228_m1 |
| mouse | interleukin 6 | *il6* | Mm00446190_m1 |
| mouse | nitric oxide synthase 2, inducible | *nos2* | Mm00440502_m1 |
| mouse | glyceraldehyde-3-phosphate dehydrogenase | *gapdh* | Mm99999915_g1 |

**Tissue processing**

Organs (heart, aorta (brachiocephalic tree, arch, thoracic and abdominal), spleen, liver, kidney, lung, muscle tissue (quadriceps), intestinal tissue) were harvested after in situ perfusion-fixation with 10 mL of PBS and 10 mL of cold 2% paraformaldehyde in PBS. Organs were placed in 2% paraformaldehyde in PBS for 2 h at 4 °C. Blood was collected into sodium citrate (3.2%) via inferior vena cava, and then plasma supernatant separated (14,000 rpm, 10 min, 4°C) and stored at -80°C until use. The heart was processed for sectioning of the aortic root as previously described[1]. Briefly, the heart was cut across in a diagonal slice from the left to right atrium and the atria placed in 30% sucrose overnight at 4 °C. Atria were quick-frozen in O.C.T. (4583; Tissue-Tek, Torrance, CA) and stored at −80 °C. 10 μm sections were cut throughout the entire aortic root and stored at -80°C. The other organs were placed in PBS with 0.05% sodium azide and stored at 4°C.

**Immunofluorescence**

Immunofluorescent (IF) staining for aortic root sections occurred following permeabilization with 0.3% Triton X-100 (X100; Sigma-Aldrich) and then were stained with 1:400 anti-CD11b antibody (NB110-89474; fisher scientific) in IHC-Tek diluent (1W-1000; IHC World, Woodstock, MD) for 1 h followed by 1:1000 Alexa Fluor 555 goat anti-rabbit IgG (A21429, Invitrogen, Carlsbad, CA) in PBS for 1 h; counterstain with 0.6 μM DAPI (D3571; Invitrogen) diluted 1:500 in PBS for 5 min was performed with all immunofluorescent staining. ProLong Gold Antifade Reagent (P36930, Thermo-Fisher Scientific) was used for mounting coverslips. Tissue sections were imaged via fluorescence microscopy using an X-Cite® 120 LED Boost (Excelitas Technologies) coupled with a Zeiss Axio Imager A.2 with either a Zeiss Objective Plan-Apochromat 5x/0.16 M27 (FWD=12.1mm), or a Zeiss Objective Plan-Apochromat 20x/0.8 (WD=0.55mm). Fluorescence microscopy images were captured with a Zeiss AxioCam HRm camera and a Zeiss 60N-C 1" 1.0X camera adapter, and observed digitally with AxioVision SE64 Rel. 4.9.1 software. Exposure times and laser strength were fixed when imaging all samples. All staining was quantified using ImageJ software (https://imagej.nih.gov/ij/).

**Oil Red O staining**

Tissue sections were fixed for 10 min with 10% neutral buffered formalin. They were then washed with water for 10 minutes,



Please do not adjust margins

rinsed with 70% ethyl alcohol for 5 mins. They were then stained for 15 min with freshly prepared Oil Red O working solution (Oil red O solution, Sigma Aldrich, O1391-500mL, dilute 3:2 in deionized $H_2O$). Stained slides were then rinsed with 70% ethyl alcohol (5 min), water (30 sec), and stained with hematoxylin (3 min), washed with $H_2O$ (2 min), then 0.5% acid alcohol solution (5 seconds; 5mL conc. HCl, 1000mL 70% ethanol), then $H_2O$ (1 min), then 0.5% ammonia water solution (10 sec; 5mL conc. Ammonium hydroxide, 1000 mL water), then finally washed with water. Slides were allowed to dry and then coverslipped and mounted with Aquamount.

#### Supplementary Results

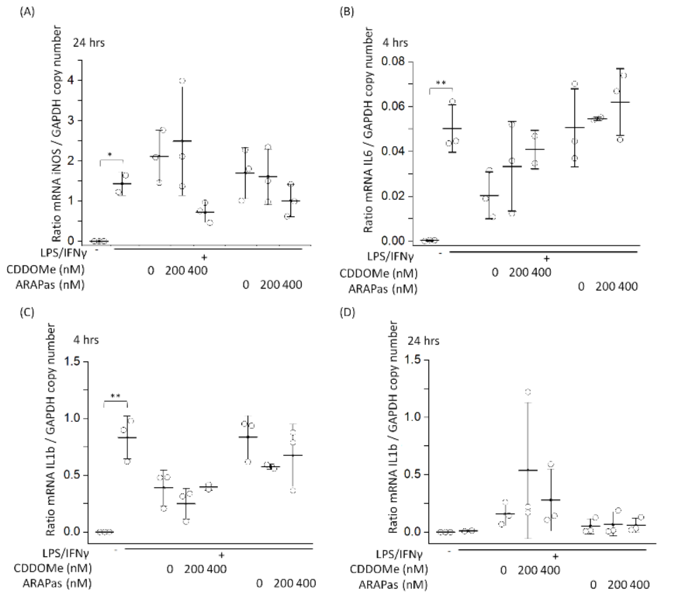


**Supplementary Figure 1. CDDOMe-ARAPas affect inflammatory signaling in classically stimulated macrophages.** RAW 264.7 macrophages were classically stimulated with IFNλ (10ng/mL, 7 hr) followed by treatment with LPS (100 ng/mL, 4-24 hrs) in the presence or absence of treatments (CDDOMe or CDODMe-ARAPas, 200-400nM) and their respective vehicles. The cells were then collected and scraped into sterile HBSS and pelleted for RNA extraction. **(A) Activation of iNOS mRNA expression following 24 hr incubation. (B) IL6 mRNA expression following 4 hr incubation. (C) IL1-β mRNA expression following 4 hr incubation and (D) 24 hr incubation.** Data represents the mean of N = 3-4 independent biological replicates, ± 1 S.D.

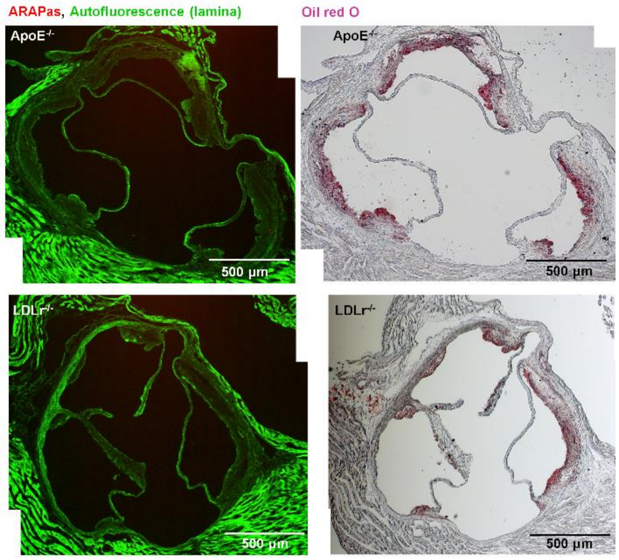


**Supplementary Figure 2. Representative images of aortic sinus of vehicle-injected animals.** LDLr$^{-/-}$ and ApoE$^{-/-}$ mice (15 wks old) were injected with PBS (5mL/kg) and their organs excised. The aortic sinus was cyro-sectioned (10 µm) and then visualized to detect DiD fluorescence. The same sections were then also stained for lipid to confirm atherosclerotic plaque presence with Oil red O staining.

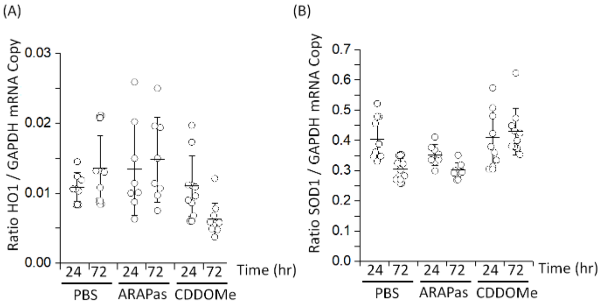


**Supplementary Figure 3. Expression of mRNA by Nrf2-regulated genes *in vivo*.** HO1 and SOD1 mRNA expression was recorded in aortic arch homogenates of high fat diet fed LDLr$^{-/-}$ mice. 4-6 wk old LDLr$^{-/-}$ mice were high fat diet fed for 8 weeks (till 12-14 wks old) and then either injected intravenously with 1mg/kg of CDDOMe encapsulated in ARAPas or 10mM PBS, or intraperitoneally with 1mg/kg CDDOme in DMSO. Aortic arches were collected at either 24 hr or 72 hr post injection of CDDOMe-ARAPas, 10 mM PBS, or CDDOme. **(A) HO1 mRNA expression in LDLr$^{-/-}$ aortic arch homegenates.** Data represents the mean of 8-10 independent biological replicates for each condition, ± 1 S.D. A 2-way ANOVA was conducted to determine the effect of treatment with either PBS, CDDOme or CDDOMe-ARAPas and time upon HO1 mRNA expression normalized to GAPDH copy number. **(B) SOD1 mRNA expression in LDLr$^{-/-}$ aortic arch homegenates.** Data represents the mean of 8-10 independent biological replicates for each condition, ± 1 S.D. A 2-way ANOVA was conducted to determine the effect of treatment with either PBS, CDDOme or CDDOMe-ARAPas and time upon SOD1 mRNA expression normalized to GAPDH copy number.


This journal is © The Royal Society of Chemistry 20xx

### ELECTRONIC SUPPLEMENTARY INFORMATION

**Supplementary Table 2. Plasma chemistry**

| Genotype | Treatment | ALB (g/dL) | ALT (U/L) | AST (U/L) | BUN (mg/dL) | Creatinine (mg/dL) | Cholestrol (mg/dL) | Triglyceride (mg/dL) | LDL (mg/dL) |
|---|---|---|---|---|---|---|---|---|---|
| ApoE$^{-/-}$ | Vehicle | 2.2 ± 0.2 | 36 ± 18 | 52 ± 9 | 22 ± 3 | 0.76 ± 0.38 | 745 ± 266 | 115 ± 88 | 242 ± 49 |
| | CDDOMe-ARAPas | 2.0 ± 0.2 | 29 ± 26 | 43 ± 14 | 15 ± 2 | 1.26 ± 0.61 | 573 ± 351 | 104 ± 54 | 247 ± 19 |
| LDLr$^{-/-}$ | Vehicle | 2.4 ± 0.2 | **87 ± 49** | **80 ± 26** | 18 ± 1 | 0.20 ± 0.07 | 1111 ± 448 | 395 ± 242 | 452 ± 191 |
| | CDDOMe-ARAPas | 2.4 ± 0.2 | **175 ± 82** | **158 ± 41** | 15 ± 2 | 0.17 ± 0.10 | 1047 ± 424 | 370 ± 172 | 389 ± 191 |
| C57Bl/6 | Vehicle | 2.2 ± 0.9 | 21 ± 4 | 32 ± 4 | 19 ± 9 | 0.25 ± 0.06 | 117 ± 8 | 84 ± 2 | 8 ± 14 |

*Data is represented as the mean of N = 5 animals per group, ± 1 standard deviation*

ELECTRONIC SUPPLEMENTARY INFORMATION
